# In-depth analysis of the tear fluid glycoproteome reveals diverse lacritin glycosylation and spliceoforms

**DOI:** 10.1101/2025.06.13.659589

**Authors:** Vincent Chang, Keira E. Mahoney, Isaac Lian, Ryan Chen, Nara Chung, Tor Paaske Utheim, Niclas G. Karlsson, Stacy A. Malaker

## Abstract

Tear fluid comprises a diverse group of extracellular glycoproteins which are critical for ocular homeostasis. Within the tear fluid glycoproteome, lacritin is highly expressed and plays a key role in immune response, tear secretion, and antimicrobial activity. Importantly, glycosylation constitutes over 50% of lactritin’s molecular weight. However, despite this fact, nothing is known about the specific glycan structures on lacritin and how they influence its protein folding, function, or downstream biological processes. Similarly, it remains completely unknown whether alterations to lacritin glycans are correlated with ocular pathologies. To address this gap in knowledge, we harnessed mass spectrometry (MS) to conduct the first O-glycoproteomic study of tear fluid. Here, we report unprecedented coverage of lacritin glycosylation, detailing 19 O-glycosites bearing a myriad of glycan structures. Further, we leveraged Alphafold 3.0 and GlycoShape to visualize the impact of these glycans on its structure, demonstrating that O-glycosylation renders the protein backbone rigid and extended. Surprisingly, we also detected protein-level evidence of two lacritin spliceoforms, representing the first observation of these isoforms by MS. Simultaneously, we describe the most comprehensive characterization of the tear fluid glycoproteome to date, elucidating the glycosylation profile of Immunoglobulin A (IgA), lactoferrin, and other glycoproteins with demonstrated clinical relevance as diagnostic biomarkers. Overall, this study lays critical groundwork for future biochemical investigation of tear fluid glycoproteins and their application as diagnostic or therapeutic tools for ocular diseases.

## Introduction

Tears are a complex extracellular fluid comprised of lipids, electrolytes, small molecule metabolites, and glycoproteins.^1^ Together, these biomolecules play key roles in maintaining ocular homeostasis and epithelial health. In dry eye disease (DED), tissue integrity and corneal maintenance is disrupted, resulting in chronic pain, impaired vision, and lasting corneal dysfunction.^2–4^ Currently, DED affects 5-50% of the world’s population,^5,6^ yet limited molecular diagnostics exist while therapies only provide modest relief to patient symptoms. Within the tear fluid glycoproteome, lacritin stands out as a glycoprotein essential for tear secretion, eye lubrication, antimicrobial activity, and immune modulation.^7,8^ Given its critical role at the ocular surface, it is perhaps unsurprising that lacritin expression levels change significantly in DED;^9–12^ as such, it is currently being investigated as a diagnostic biomarker. Beyond its diagnostic capacity, lactritin bears a peptide epitope (residues 114 to 138) that has recently been developed into Lacripep^TM^, a first-in-class therapeutic candidate for DED which aims to reverse pathological outcomes of dry eye.^13,14^

These advances in lacritin-based diagnostics and therapeutics are largely driven by key biochemical discoveries made by the Laurie group over the past 3 decades. Previously, they demonstrated that the biological activity of lactritin is mediated by its ability to bind syndecan-1, which initiates calcium signaling and mitogenic activity toward epithelial repair.^15,16^ Importantly, signaling through the lacritin-syndecan-1 axis is established through noncovalent interactions between the C-terminal alpha helix of lacritin (residues 114 to 138) and the N-terminal residues (1 to 51) of syndecan-1. Interestingly, lacritin is also known to form dimers, trimers, and multimers through transglutaminase (TGM2)-mediated crosslinking, where Lys101 and Lys103 act as donor residues and Gln125 acts as an acceptor residue.^17^ Although the role of lacritin multimers is unclear, they bind syndecan-1 less effectively than monomeric lacritin, suggesting that multimerization impairs syndecan-1 binding and thus downstream lacritin effector function. In addition, elevated levels of TGM2 are implicated in ocular diseases such as DED^18^ and glaucoma^19^, where negative regulation of monomeric lacritin by TGM2 could be an underlying mechanism.

Despite over 30 years of literature on lacritin, the glycan structures which decorate its protein backbone remain completely unknown. Glycans comprise more than half of its molecular weight;^20^ however, their impact on lacritin structure, function, and protein-protein interactions remains entirely unexplored. Most commonly, glycans attach to asparagine (N-linked) and serine/threonine residues (O-linked) in a non-templated fashion. Crucially, glycans mediate a host of biochemical processes, including protein folding, cellular adhesion, and immune signaling.^21,22^ Additionally, dysregulation of glycosylation is concomitant with a wide range of diseases, including ocular pathologies such as diabetic retinopathy^23^ and atopic/vernal keratoconjunctivitis (AKC and VKC).^24^ Overall, molecular-level insight into the glycosylation landscape of lacritin will have significant implications for unraveling biology dictated by its glycans and its potential dysregulation in various diseases. Furthermore, site-specific elucidation of lacritin glycoepitopes could aid in the development of glycosylation-centric diagnostic tools and therapeutic modalities for ocular diseases where lacritin expression and glycosylation are dysregulated.

Glycoproteomics, or the systems-level study of glycoproteins using MS, has emerged as the premier analytical strategy to reveal key glycosylation signatures in complex biological samples.^21,22^ As lacritin is predicted to be heavily O-glycosylated, we reasoned that a targeted O-glycoproteomic analysis of tear fluid could elucidate its endogenous glycosylation landscape. Currently, however, glycoproteomic profiling of tear fluid presents an enormous analytical challenge due to the low concentration of proteins in tears, low volume of sample collection, and sheer complexity of the glycoproteome. This is further highlighted by the fact that the two existing N-glycoproteomic studies required very large sample input, thus necessitating patient samples to be pooled for analysis.^25,26^ Additionally, these studies employed the endoglycosidase PNGaseF for analysis. This enzyme removes endogenously expressed N-glycans, leaving a deaminated scar in its place, and is often used to site-localize where N-glycans were present. That said, this technique cannot reveal which glycans are found at individual residues, thus leaving critical information unstudied. Finally, O-glycoproteomic studies have never been performed using tear fluid, further highlighting the difficulty in studying this type of sample.

To address this gap in knowledge, we sought to implement intact glycoproteomics workflows capable of characterizing tear fluid glycoproteins from single patients. Previously, our lab developed two O-glycoprotein enrichment strategies that employ a class of recently introduced enzymes termed “mucinases”. Briefly, mucinases are a subclass of proteases which specialize in digestion of proteins bearing densely O-glycosylated domains (herein referred to as mucin domains) to generate glycopeptides amenable to MS analysis.^27–29^ In prior studies, we reported the use of two mucinases, *Serratia marcescens* Enhancin (SmE)^29^ and secreted protease of C1 esterase inhibitor (StcE)^30^, to aid in O-glycoproteomic analysis. In our first enrichment method, an O-glycosylation-focused filter-aided sample preparation (GlycoFASP) workflow generates abundant O-glycopeptides that can be separated from non-glycosylated proteins using a molecular weight cut off (MWCO) filter.^31^ The other preparation method employs an inactive point mutant of StcE (StcE^E447D^), which binds mucins without cleaving them. Here, StcE^E447D^ is immobilized onto a solid support to selectively enrich mucin-domain glycoproteins from complex biological samples.^32^

Leveraging these methods, we describe unprecedented coverage of the tear fluid glycoproteome, highlighting diverse lacritin glycoforms bearing highly heterogeneous glycan structures. Building on this, we performed protein modeling with Alphafold 3.0 and GlycoShape to visualize the chemical influence imparted by glycans on lacritin’s secondary structure. Furthermore, we report protein-level evidence of two lacritin spliceoforms, which may help to explain the diversity of lacritin glycoforms. Finally, beyond lacritin, we elucidate the glycosylation profile of IgA and other glycoproteins known to play key roles at the ocular surface. Overall, the findings from this study lay the critical groundwork for future investigation of lacritin and other tear fluid glycoproteins in ocular diseases such as DED.

## Results and Discussion

### Development of a new glycoproteomic workflow for tear fluid

Here, we adapted our GlycoFASP strategy^31^ to analyze tear fluid O-glycoproteins derived from Schrimer tear strips (**Fig. 1A**), where a typical collection yields up to 80 µg of protein.^33^ Briefly, our method employs a protein extraction protocol where reduction with Dithiothreitol (DTT) is performed directly on the Schirmer strip in the presence of a chaotropic agent (**Fig. 1A**). Following extraction, proteins are alkylated with iodoacetamide (IAA) and loaded onto a 50 kDa filter for several rounds of washes. Finally, an O-glycoprotease (mucinase SmE) is added for on-filter protein digestion and glycopeptides passed through the filter are collected for desalting and subsequent LC-MS/MS analysis. When compared to an unenriched control (representative of proteomic sample processing methods currently reported in the literature), GlycoFASP boasted an ∼4 fold increase in O-glycopeptide (57 to 233) identifications (IDs), and an ∼2 fold increase in N-glycopeptide IDs (23 to 53) (**Fig. 1B Table S1**).

**Figure 1.**
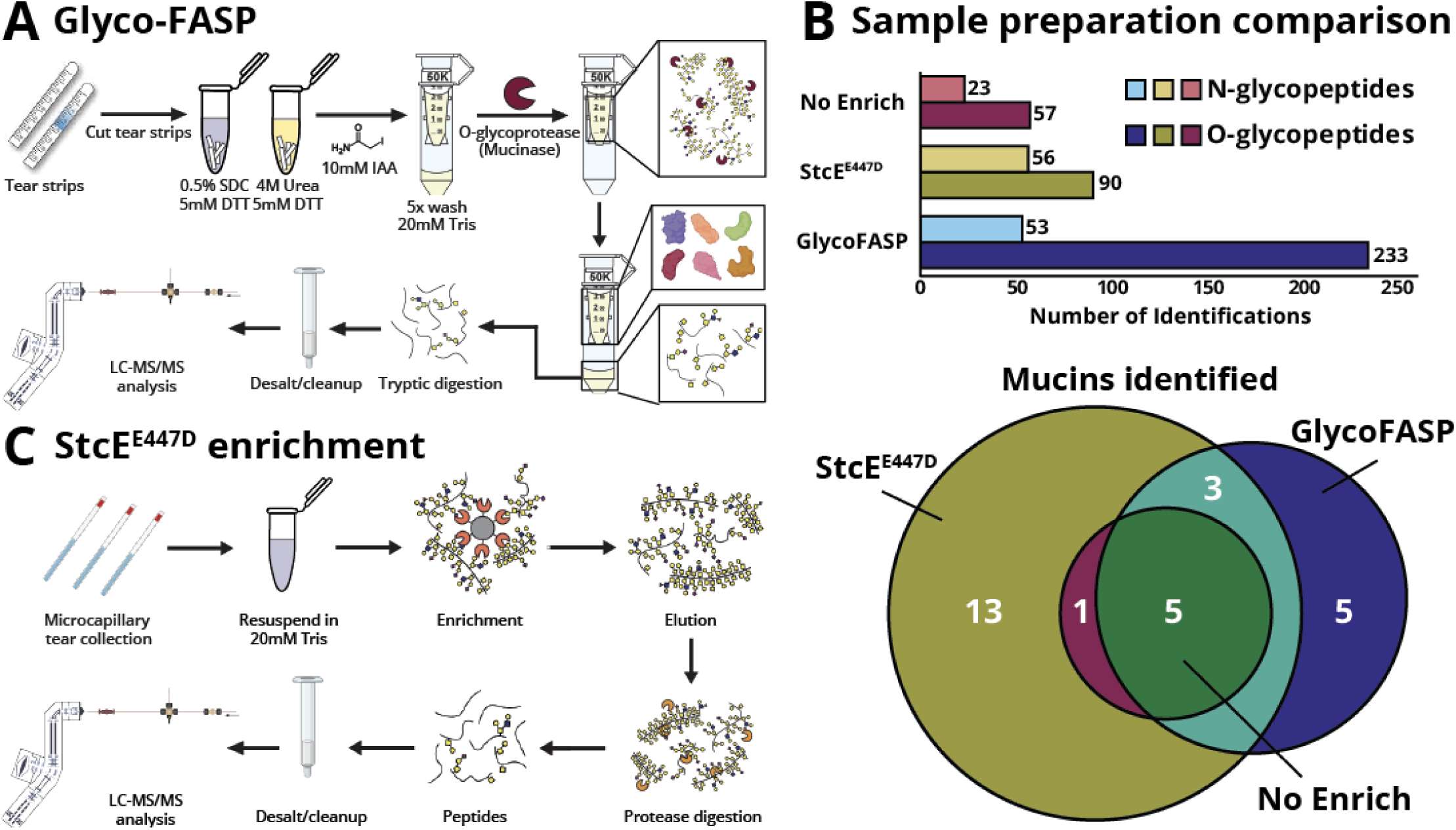
Glycoproteomic enrichment methods for tear fluid. **(A)** GlycoFASP strategy performed on tear fluid proteins extracted from Schirmer tear strips. Proteins are reduced and alkylated following extraction and loaded onto a 50KDa MWCO filter before protease digestion and subsequent MS analysis. **(B)** StcE^E447D^, an inactive mucinase, is conjugated onto a solid support and used to enrich mucin-domain O-glycoproteins from tears collected by capillaries. Following enrichment and elution, proteins are digested with proteases before MS analysis. **(C)** Sample preparation comparison for N- and O-glycopeptide IDs (top) and mucin IDs (bottom). In total, 22 mucins were IDed with StcE^E447D^ enrichment, 6 mucins with no enrichment, and 13 mucins with GlycoFASP.

While GlycoFASP enabled us to generate a high abundance of O-glycopeptides, the number of mucin-domain glycoprotein IDs (13 in total) was fewer than we initially anticipated given that abundant mucin expression is essential for ocular homeostasis.^34^ To improve global identification of mucin-domain O-glycoproteins, we opted for an alternative enrichment workflow which immobilizes StcE^E447D^ onto a solid support, thus allowing us to selectively enrich mucin-domain glycoproteins from complex samples (**Fig. 1C**).^32^ As previously reported, this method requires higher sample input relative to GlycoFASP (300 µg vs 50 µg of protein). We therefore reasoned that tears derived from microcapillary sampling, where collections from the same patient throughout the day could yield sufficient protein quantities, would be suitable for this enrichment method. Compared to an unenriched control, StcE^E447D^ enrichment similarly resulted in higher O-glycopeptide IDs (57 to 90), and N-glycopeptide IDs (23 to 56). Notably, however, mucin glycoprotein IDs more than tripled (6 to 22) relative to an unenriched control (**Fig. 1B, Table S1**). Taken together, these results highlight the versatility of our workflows in capturing the glycosylation landscape of tear fluid glycoproteins.

### Glycosylation landscape of lacritin

Elucidating the site-specific glycan structures which decorate lacritin has significant implications for furthering our molecular-level understanding of lacritin’s biological roles. With this in mind, we harnessed our glycoproteomic workflow to comprehensively map the glycosylation landscape of endogenous lacritin (**Fig. 2A**). Prior to this study, 14 O-glycosites were predicted based on its protein sequence using NetOGlyc 4.0,^20^ which leverages a neural network to predict sites of O-linked glycosylation based on experimental evidence. Notably, we were able to detect all 14 predicted sites as well as an additional 5 O-glycosites that were not predicted by NetOGlyc. Many of these were modified by a range of different glycan structures. For instance, we observed the Tn antigen (GalNAcα1-Ser/Thr; 89% relative abundance), sialylated core 1 (7.35%), type 3 H-antigen (Fuc α1-2-Galβ1-3GalNAcα1-Ser/Thr; 1.89%), and sialylated and fucosylated core 2 (0.30%) O-glycan structures, which were quantified by label free quantitation (LFQ) using area under the curve (AUC) intensities of extracted ion chromatograms (XICs) (**Fig. 2B**). Overall, the O-glycans we site-localized onto lacritin corroborate many of the O-glycan structures observed in previous glycomic studies of basal tears,^35,36^ including the sialyl Lewis x epitope and the H blood group antigen. Additionally, we note that regions of lacritin known to be important for cell adhesion^37,38^ and syndecan binding^15,16,39^ contain glycosylated Ser, Thr, or Asn residues. As such, it is likely that distinct lacritin glycoforms have unique roles in tear fluid and future studies to elucidate their specific functions are warranted.

**Figure 2.**
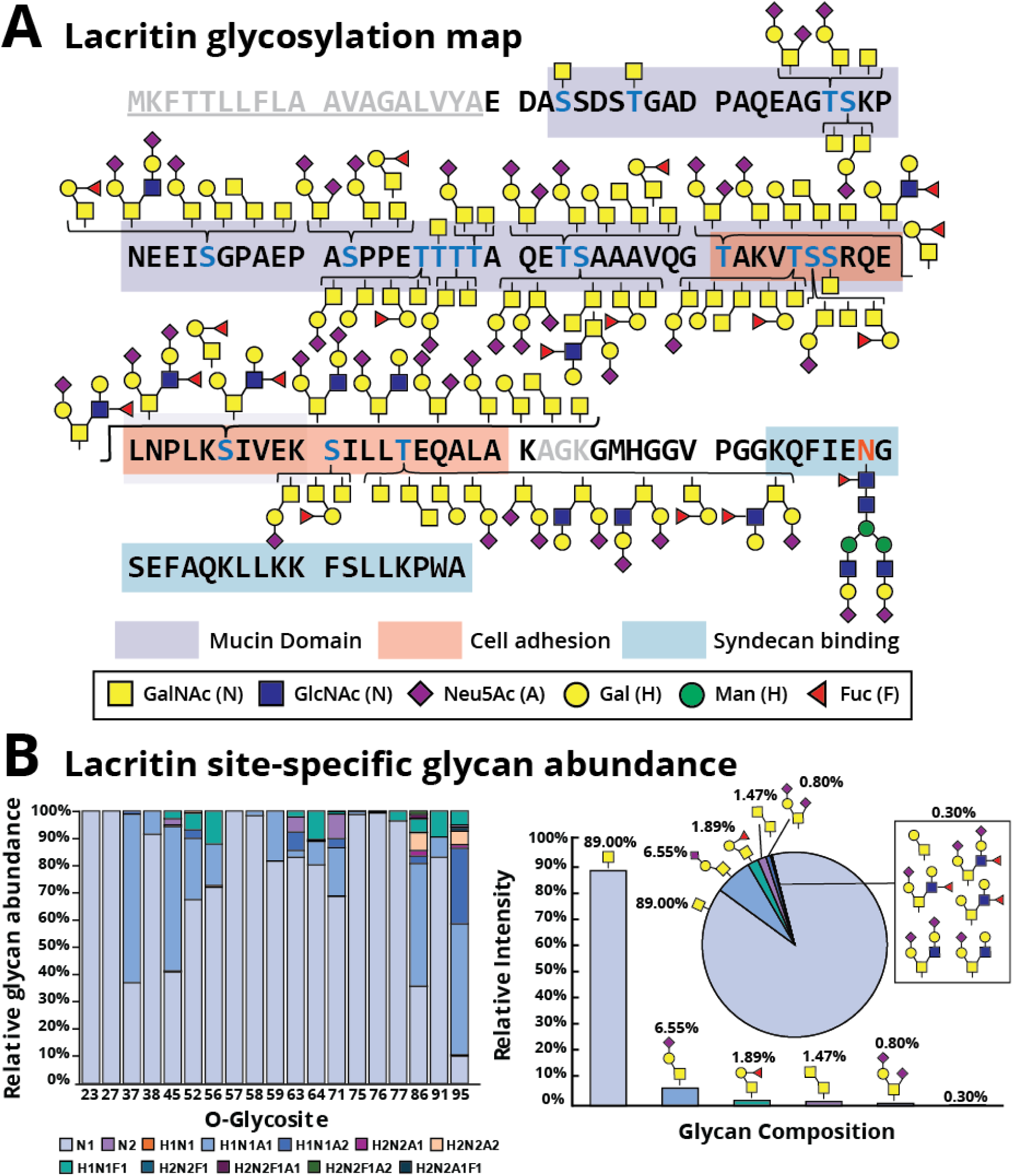
Glycosylation landscape of lacritin. **(A)** Full sequence of the canonical isoform of lacritin. The signal sequence is underlined and shaded in gray, localized O-glycosites are colored in blue, and the N-glycosite is colored in orange. Regions important for cell adhesion (orange), syndecan binding (blue), and representing the mucin domain (purple) are also shown. **(B)** Site-specific quantification of lacritin at each glycosite (left). Relative O-glycan abundances across all O-glycosites. Quantification was performed using label free quantitation (LFQ) and area under the curve (AUC) values from extracted ion chromatograms (XICs) of all identified glycopeptides.

### Lacritin protein modeling with Glycoshape and AlphaFold 3.0

It is possible that lacritin glycans mediate its protein-protein interactions (PPIs) given their proximity to key crosslinking amino acid residues and their ability to dramatically affect protein structure and folding. To address the influence of glycans on the secondary structure of lacritin, we sought to convert our 2D lacritin glycomap into a glycosylated 3D protein structure. To do so, we first calculated the most abundant glycan structure at each glycosite (**Fig. 2B**). Interestingly, the predominant glycan structure was the Tn antigen for all glycosites except Thr37, Ser 45, Ser 86, and Thr95. Next, we leveraged a recent glycoprotein modeling software, GlycoShape^40^, which can site-specifically incorporate glycans onto the protein backbone. Importantly, GlycoShape encompasses a wide repertoire of available glycan structures, which made it an attractive starting point for glycan visualization. To date, the best structure prediction of lacritin originates from AlphaFold 3.0 (**Fig. 3A**). Comparatively, the GlycoShape-predicted structure (**Fig. 3B**) highlights the large steric contribution imparted by lacritin N- and O-glycans. While this was a promising start to revise the proposed structure of lacritin, we realized that this model did not significantly change the predicted fold of the protein. Indeed, GlycoShape was primarily designed for visualization of N-glycosite occupancy where the fold of the protein is already structured. For mucin domains which are predicted to be intrinsically disordered, GlycoShape can sample 3D space accessibility at O-glycosites to evaluate steric clashes; however, the base protein fold predicted by Alphafold remains unchanged. Given the density of O-glycosylation on the lacritin protein backbone, we expected that it might adopt the typical “bottle-brush-like” structure characteristic of mucin-domain glycoproteins.^29,41,42^ Inspired by the new post-translation modification (PTM) feature of AlphaFold 3.0, we repeated the process of adding N- and O-glycans to the backbone of lactrin and, as seen in **Fig. 3C**, we observed a dramatic change in the predicted protein fold (or lack thereof, with respect to the mucin domain). Overall, this supports that lactritin glycosylation is predicted to significantly change the secondary structure of lacritin.

**Figure 3.**
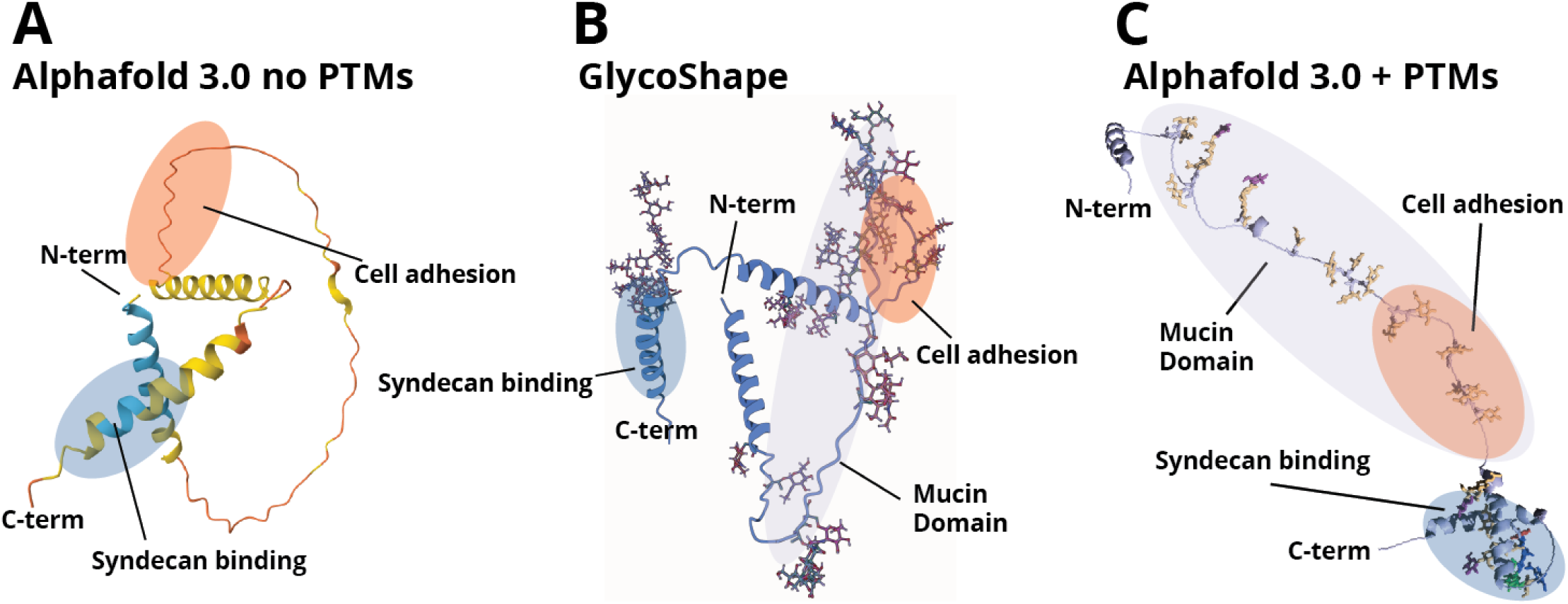
Protein modeling of lacritin. **(A)** Current predicted structure of lacritin with AlphaFold 3.0 and no PTMs. **(B)** GlycoShape predicted structure of lacritin. **(C)** AlphaFold 3.0 structure with PTMs. O-glycans used for GlycoShape and AlphaFold 3.0 are based on only the most abundant O-glycan structure at each glycosite.

In both the protein models we investigated, glycans were found within regions of lacritin that are important for PPIs. Prior studies by the Laurie group on the lacritin-syndecan-1 axis revealed that the non-glycosylated C-terminal helix of lacritin was sufficient to bind syndecan-1 and induce downstream signaling.^15,16,43^ Given that we observed an occupied N-glycosite within this helix, we first wanted to visualize the steric contribution of this glycan with GlycoShape (**Fig. S1**). As expected, the size of the glycan within this helix is significant and would likely affect or prevent syndecan-1 binding. We therefore aimed to quantify the relative abundance of this glycoform in comparison to its nonglycosylated counterpart (**Fig. S2**). Interestingly, the relative abundance of the unmodified peptide was significantly higher, accounting for 99.3% of the relative intensity while the N-glycosylated peptide accounted for only 0.7%. Based on the low abundance and large steric contribution of this N-glycan, it is possible that monomeric populations of lacritin which engage syndecan-1 do not bear this glycoepitope. The presence of this N-glycan was previously unreported, and its role therefore remains unclear. It may function to affect lacritin C-terminal proteolytic processing, transglutaminase-mediated lacritin multimerization via Gln125, or engage another protein entirely. Future biochemical studies with an N-glycosylated helix of lacritin will be necessary to determine the extent to which the N-glycan mediates or abrogates binding to syndecan-1.

### Identification of lacritin isoforms A, C, and D

Based on the vast diversity of lacritin glycoforms we observed, we asked whether splice variants were also detectable and potentially contributing to the heterogeneous population of lacritin. Indeed, several prior studies have demonstrated that protein splice variants could harbor distinct glycosylation signatures.^44–46^ Most recently, lubricin, a mucin-domain glycoprotein important for joint lubrication, was reported to exhibit multiple splice variants which displayed isoform-specific O-glycan profiles.^47^ Here, we reasoned that lacritin splice variants may also be present in our analysis. To date, outside of the canonical isoform A, three additional mRNA splice variants of lacritin (isoforms B,C,D) have been identified by transcriptomics albeit in relatively low abundance.^10,20^ Of these isoforms, only isoform C has been detected at the protein-level with an isoform-specific antibody^48^, though it remains to be validated by MS-based proteomics. Interestingly, isoform D is not predicted to be translated based on its short sequence length and lack of a signal peptide, though the entirety of its O-glycosylated mucin domain is encoded.

To investigate the presence of different lacritin splice variants, we first determined tryptic peptide sequences unique to each isoform (**Fig. 4A**). The peptides SILLTEQALAK, QELNPLSK, and QFIESECIPR provided evidence for spliceoforms A, C, and D, respectively. Notably, the tryptic peptide QELNPLKQALAK (containing one missed cleavage) from isoform B is distinct from A and C, though this sequence is shared with isoform D. Regardless, this peptide was not observed in this study, so we cannot comment on the presence of isoform B. After determining these sequences, we extracted the ion chromatograms (XICs) for the peptide QLENPLSK (isoform C) at *m/z* 469.759 and QFIESECIPR (isoform D) at *m/z* 639.812; we then confirmed the sequence assignment using MS2 (**Fig. 4B** and **C**). For isoform A, we identified both glycosylated and unmodified versions of SIVEKSILLTE, which was quantified using LFQ of AUC intensities from XICs of each glycoform (**Fig. S3**). Quantification revealed that the unmodified isoform A peptide had significantly higher relative abundance (6.97*10^10^ AUC intensity) compared to the three other glycoforms, which exhibited either a Tn antigen or a disialylated core 1 O-glycan (**Fig. S3**). Furthermore, the intensity of the isoform A specific peptide (6.97*10^10^) was much higher compared to that of isoforms C (1.36*10^7^) and D (1.48*10^9^).

**Figure 4.**
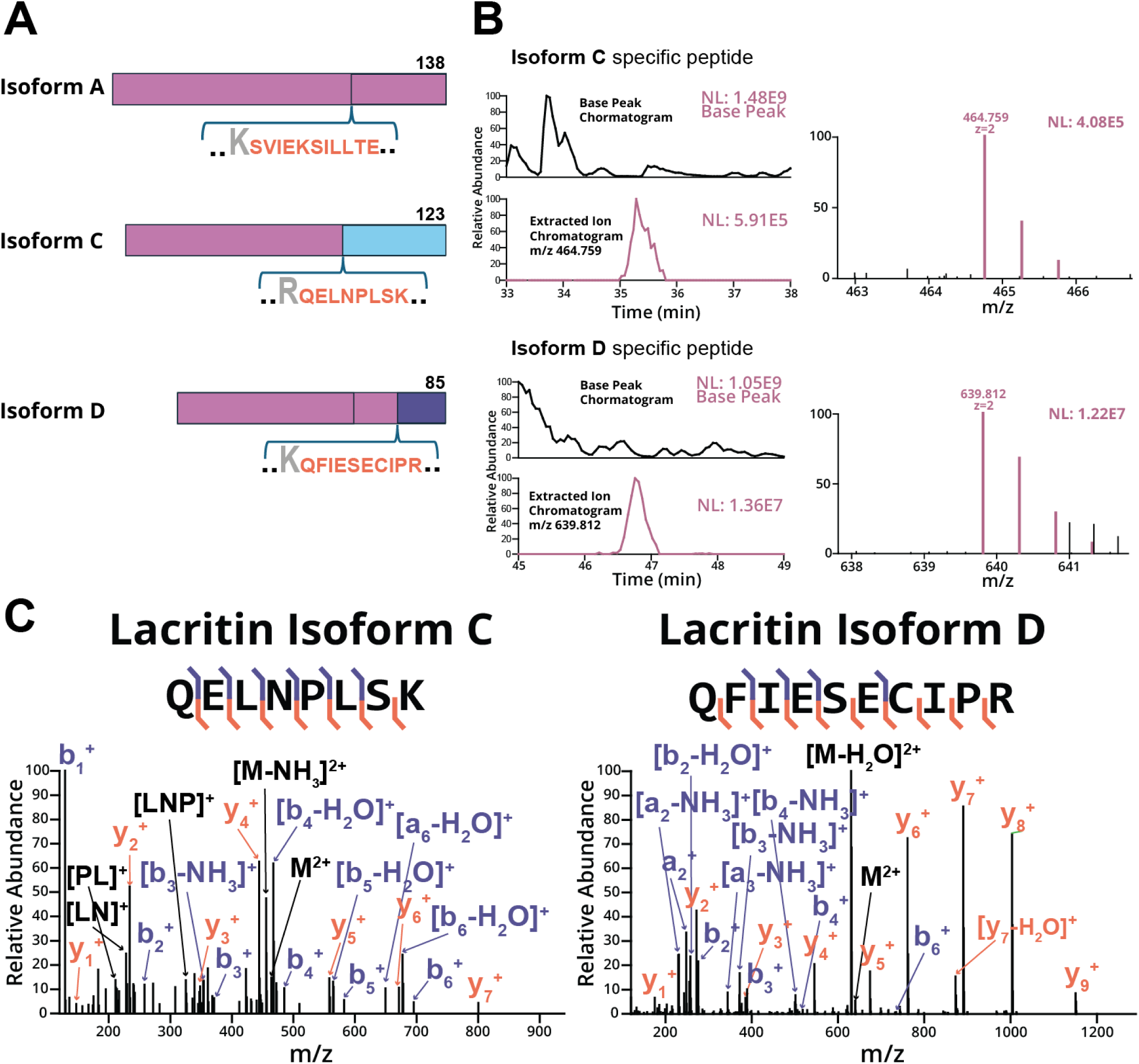
lacritin isoforms from alternative splicing. **(A)** Visual representation of lacritin isoform and sequence-specific peptides. **(B)** XIC and base peak chromatogram for the isoform C-specific peptide (m/z=464.759, z=2, retention time 33-38 minutes) and the isoform D-specific peptide (m/z 639.812, z=2, retention time 45-49 minutes). The MS1 of the monoisotopic precursor and its isotopes are shown (right) along with other co-isolated species. **(C)** Annotated MS2 spectra for isoform-specific peptides C and D.

Discovery of lacritin isoforms C and D by MS has important biological implications for contextualizing the multifaceted role of lacritin at the ocular surface. Over three decades of literature on lacritin has largely focused on isoform A, leaving the function of other isoforms completely uninvestigated. Most surprisingly, we described the existence of isoform D circulating in tear fluid, an 85 amino acid splice variant without an annotated signal sequence and predicted not to be translated. To better understand the structure of these isoforms, we again performed AlphaFold modeling and showed that both isoform C and D are predicted to adopt an alpha helical structure within their isoform-specific sequences (residues 83 to 101 and 53 to 63, respectively). A previous study by Zhang et al. on isoform C corroborates an ordered structure around this region which is unable to bind syndecan-1, leaving its current function unknown.^39^ Further analysis of these two helices revealed a difference in length, amino acid identity, and hydrophobicity compared to the C-terminal alpha helix observed in isoform A (**Fig. S4**). Taken together, this would support the previous observations on isoform C made by Zhang et al., though it remains unclear whether isoform D would be able to engage syndecan-1 without further biochemical characterization. Notably, isoform D contains Gln and Lys acceptor residues for TGM2-mediated crosslinking to form multimers. Given that isoform A and C have previously been shown to form multimers, isoform D may also multimerize.

Lastly, we performed multiple sequence alignment (MSA) analysis which revealed that the evolutionary conservation of these isoforms dates back only to a few primates, while the densely O-glycosylated region of lacritin seems to remain conserved across a wider range of organisms (**Fig. S5**). The evolutionary persistence of the mucin domain within lacritin suggests a key biological role associated with this densely O-glycosylated region. It is possible that the mucin domain serves a protective role against proteases^49,50^, acts as a ligand for galectin-3 in tear fluid^51^, or mediates lacritin crosslinking, which can be elucidated in future studies.

### Uncovering the tear fluid glycoproteome

Beyond lacritin, we performed the most in-depth analysis of the tear fluid glycoproteome to date. Here, we report a range of site-localized N- and O-glycans, including high mannose, hybrid, and complex (sialylated, fucosylated, and sialofucosylated) N-glycans and Tn antigen, core 1, and core 2 O-glycans **(Fig. 5A)**. While this corroborates previous N- and O-glycomic studies on tear fluid^23,24,52,53^, we note that site-specificity and protein identity is inherently lost in glycomics experiments. Therefore, we aimed to elucidate the site-specific glycosylation landscape of O-glycoproteins identified with GlycoFASP, which allowed us to observe vastly diverse O-glycosylation profiles unique to individual proteins (**Fig. 5B, Table S1**). For instance, proline-rich protein 4 (PROL4) primarily displayed the Tn antigen, while deleted in malignant brain tumors 1 (DMBT1) was predominantly sialylated. This may suggest that these proteins derive from different ocular cell types with distinct glycosyltransferase expression levels. Alternatively, the sequence of the glycoprotein itself may exhibit different propensities for glycan extension based on solvent accessibility and specific sequence motifs for extending glycosyltransferases.^54^ To better understand the function of glycoproteins captured by GlycoFASP, we performed relative quantitation (**Fig. 5C**) and GO enrichment analysis (**Fig. 5D**). Overall, the top 10 highest intensity proteins identified with this method overlap with previous proteomic analyses of tear fluid, and the biological functions of the broader glycoproteome also support previous observations.^25,55,56^

**Figure 5.**
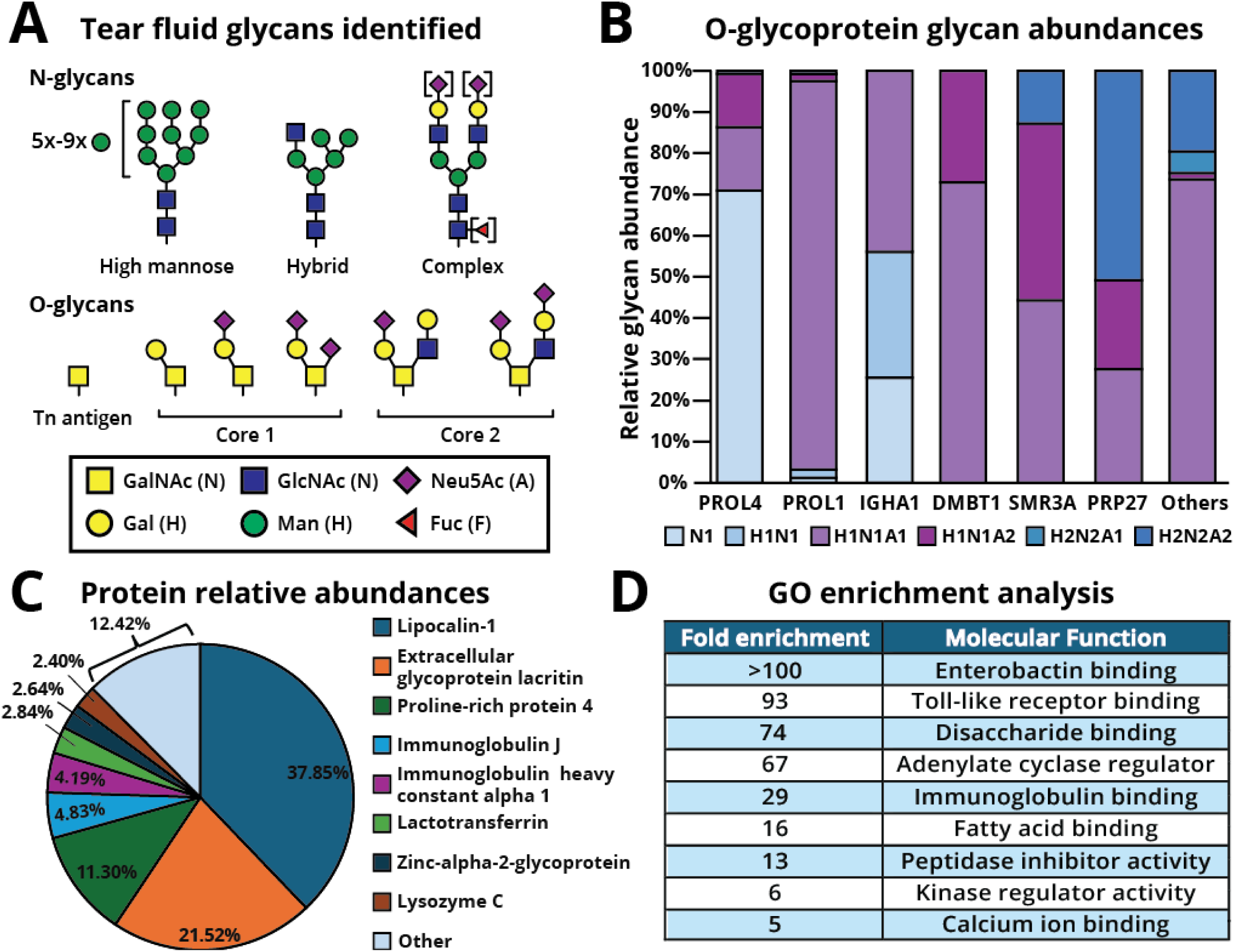
Uncovering the tear fluid glycoproteome. **(A)** N- and O-glycan structures identified in this study using GlycoFASP method (**B**) O-glycosylation landscape of different O-glycoproteins where H represents Hexose, N represents HexNAc, and A represents Neu5Ac. (**C**) Protein relative abundances from glycoFASP. (**D**) Gene ontology (GO) enrichment analysis of all glycoproteins identified from both enrichment methods.

### In-depth glycosylation characterization of secretory IgA1 and IgJ in tears

IgA1 is the most abundantly expressed immunoglobulin in the human body, where it plays an important role in mucosal immunity and protection against pathogens. Structurally, it is often found as a secreted homodimer joined together by an N-glycosylated IgJ chain (**Fig. 6A**). Given that IgA1 has documented glycan alterations in various diseases,^57,58^ we asked whether our workflow could enable deep glycoproteomic profiling of IgA1 and its glycoforms, with the goal of informing future diagnostic tools. Notably, previous work revealed that aberrant glycosylation of IgA1 plays a key role in the pathogenesis of autoimmune diseases such as IgA nephropathy (IgAN), and IgA vasculitis (IgAV).^59,60,61^ Furthermore, altered glycosylation of serum IgA1 is associated with ovarian cancer, breast cancer, colorectal cancer, and hepatitis B virus-related liver cancer.^59,62–64^ While serum IgA has been extensively investigated as a glycan-specific disease biomarker, tear fluid IgA and its specific glycoforms have yet to be elucidated, where its glycosylation could be diagnostically informative for diseases such as dry eye disease (DED). Here, we reasoned that if our workflow could capture tear fluid IgA glycoepitopes, then our enrichment strategy could be applied in the future to profile IgA glycosylation in ocular diseases where aberrant glycosylation may be prevalent.

**Figure 6.**
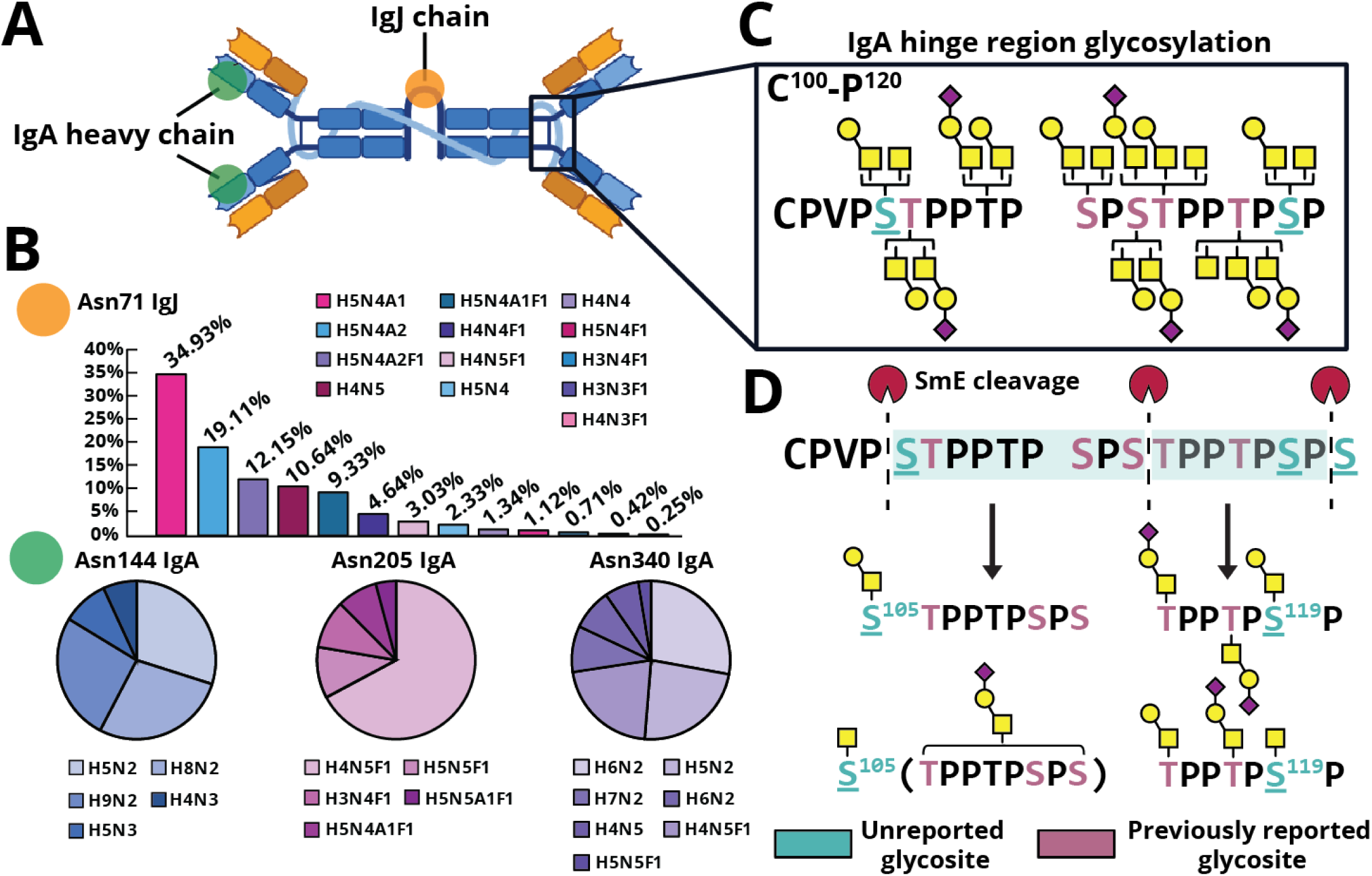
IgA and IgJ site-specific N-/O-glycan analysis. **(A)** Cartoon representation of IgA dimer held together by IgJ. **(B)** Site-specific O-glycosylation of IgA hinge region with previously unreported glycosites colored in teal and previously reported glycosites colored in pink. **(C)** Site-specific N-glycan analysis of IgA and IgJ where H represents Hexose, N represents HexNAc, F represents Fucose, and A represents Neu5Ac. **(D)** Representation of glycopeptides generated by mucinase SmE to enable glycosite localization.

Previous studies reported a total of 6 O-glycosylation sites in the IgA hinge region (HR), where an increase in the Tn antigen or sialyl Tn antigen (Neu5Acα2-6GalNAcα-Ser/Thr) and a decrease in sialyl T antigen (Galβ1-3GalNAcα1-Ser/Thr) are concomitant with IgAN.^57,65–67^ Historically, characterization of IgA1 HR glycosylation has been analytically challenging.^66,67^ This is evidenced by the fact that previous glycoproteomic studies of IgA1 are performed on highly purified IgA1, necessitating both immunoprecipitation and size-exclusion chromatography procedures on large quantities of human serum.^65,68^ In this study, we demonstrated full coverage of the IgA1 HR from just 50 µg of tear fluid, where we site-specifically localized Tn and (sialyl) T antigen O-glycan structures to all 6 of the previously reported glycosites (shown in pink) (**Fig. 6C** and **6D**). In addition to these known glycosites, we identified two previously unreported glycosites (teal), which we were able to localize via digestion with SmE (**Fig. 6D**). The glycopeptides generated by SmE are depicted with their corresponding O-glycan structures. Overall, we describe comprehensive coverage of the IgA1 hinge region without tedious purification procedures while requiring minimal sample input. This not only represents the first characterization of IgA1 glycoforms in tear fluid but also highlights the sensitivity of our workflows, which we envision can be applied to other biofluids with ease.

In addition to mapping the densely O-glycosylated HR, we performed site-specific N-glycan quantification of Asn71 on IgJ and Asn144, 205, and 340 of IgA1 (**Fig. 6B**). Interestingly, sialylated and sialofucosylated N-glycan structures were highly expressed on IgJ, while IgA1 exhibited almost no sialylation on its N-glycan structures (∼10% relative abundance on only Asn205 and no other glycosites). Further, the N-glycan structures modifying the 3 N-glycosites of IgA1 varied drastically, as evidenced by the extensive fucosylation on Asn205 when compared to Asn144 (no fucosylation) and Asn340 (<10% relative abundance of fucosylation). Finally, high mannose N-glycans seemed to predominate on Asn144 and Asn340, though complex N-glycans were also detected at these sites. Our results corroborated previous tear fluid N-glycomics studies which suggested that tear fluid N-glycans are primarily complex structures and heavily fucosylated, which was observed on Asn71 of IgJ and Asn205 of IgA1.^69^ Additionally, the identified high mannose structures on Asn144 and Asn340 of IgA1 have similarly been reported in a separate N-glycoproteomic study on serum and salivary IgA1.^68^

Beyond IgA1 and IgJ, we also investigated N-glycosylation at Asn497 on lactoferrin (**Fig. S6**), a highly abundant glycoprotein in tear fluid (making up 25% of total tear protein abundance)^70^ with known changes in expression levels in DED. Lactoferrin plays important roles at the ocular surface through its antibacterial activity, anti-inflammatory properties, and ability to promote cell proliferation. While the role of glycans on lactoferrin is not well understood, alterations to its N-glycan structures have been reported in several diseases.^71,72^ Future application of our workflow to investigate lactoferrin glycosylation could also reveal N-glycan changes which are associated with specifically ocular pathologies.

## Conclusion

This study represents the most comprehensive analysis of the site-specific glycosylation landscape of tear fluid to date. To overcome previous analytical challenges in studying tear fluid, we harnessed recently introduced mucinases and unique enrichment techniques, which enabled unprecedented coverage of the tear fluid glycoproteome. We highlight the versatility of our methods through site-specific profiling of glycosylated proteins which are established diagnostic biomarkers and key regulators of ocular homeostasis, namely, IgA1 and lacritin. Specifically, lacritin has been extensively studied over the past 30 years and is the only therapeutic for DED currently in clinical trials. Despite this, the role of the glycans which occupy lacritin and comprise more than half its molecular weight were previously a critical blind spot, until now. Leveraging our site-specific glycomap of lacritin, we employed two protein modeling programs, GlycoShape and AlphaFold 3.0, to better understand the structure of lacritin. Ultimately, we have provided new insight into the complex biology mediated by lacritin, corroborating previous observations by the Laurie group while raising new possibilities for the role of specific lacritin glycoforms and isoforms.

## Materials and Methods

### Tear fluid Sample collection

Tear fluid from an anonymous healthy volunteer was collected on Schirmer tear strips (placed onto the lower eyelid for 5 min) or by microcapillary tubes. Tears collected by microcapillary tubes were transferred into 1.5 mL Eppendorf tubes and stored at −20°C until further processing. Schirmer strips was air dried in room temperature in a dust-free environment and stored at −20°C until further processing.

### Mass Spectrometry Sample preparation GlycoFASP of tear fluid

Tear fluid proteins were directly extracted and reduced from tear strips by using either 0.5% sodium deoxycholate (SDC) in 5 mM dithiothreitol (DTT) and 20 mM Tris (65°C 1 hour), or 4 M Urea in 5 mM DTT and 20 mM Tris (23°C, 4 hours). Next, proteins were alkylated with 10 mM IAA (15 minutes in the dark at 23 °C) and subsequently washed 6 times with 20 mM Tris on a 50 kDa MWCO filter. On-filter digestion was performed with mucinase SmE (enzyme:substrate ratio of 1:20), which was allowed to react overnight at 37 °C. Glycopeptides generated from mucinase digestion can now pass through the 50 kDa filter, and are collected for subsequent tryptic digestion at a 1:50 enzyme:substrate ratio for 6 hours at 37 °C. All reactions were quenched by adding 1 µL of formic acid and diluted to a volume of 200 µL prior to desalting. Desalting was performed using 10 mg Strata-X 33 µm polymeric reversed phase SPE columns (Phenomenex). Each column was activated using 500 µL of acetonitrile (ACN) (Honeywell) followed by of 500 µL of 0.1% formic acid, 500 µL of 0.1% formic acid in 40% ACN, and equilibration with two additions of 500 µL of 0.1% formic acid. After equilibration, the samples were added to the column and rinsed twice with 200 µL of 0.1% formic acid. The columns were transferred to a 1.5 mL tube for elution by two additions of 150 µL of 0.1% formic acid in 40% ACN. The eluent was then dried using a vacuum concentrator (LabConco) prior to reconstitution in 10 µL of 0.1% formic acid. The resultant peptides were then injected onto a Dionex Ultimate3000 coupled to a Thermo Orbitrap Eclipse Tribrid mass spectrometer. We employed a higher-energy collision dissociation product-dependent electron transfer dissociation (HCD-pd-ETD) method; in some cases, we used supplemental activation in ETD (EThcD). The files were searched using Byonic, followed by manual data curation.

### Preparation of StcE^E447D^ beads

The pET28 plasmids for His-tagged SmEnhancin and StcE were kindly provided by the Bertozzi laboratory. StcE^E447D^ was conjugated to NHS-activated beads and washed 3× with 1 mL of PBS followed by the addition of the proteins, which were allowed to bind overnight at 4 °C. Free NHS-esters were capped by adding 100 mM Tris, pH 7.4, to the bead slurry for 20 min at 4 °C. BCA assays were performed after the reactions in order to determine the sufficient binding efficiency. After capping, beads were washed 3x with a high-salt buffer (20 mM Tris and 500 mM NaCl) followed by a buffer without salt (20 mM Tris).

### StcE^E447D^ enrichment of tear fluid

Alternatively, tears collected from capillary tubes (diluted to 500 µL with PBS with a final concentration of 0.5 mg/mL) were loaded onto StcE^E447D^-conjugated NHS beads, washed 3 times with 20 mM Tris and 500 mM NaCl, and eluted with 0.5% SDC. Following elution, the mucin-enriched tear fluid was subjected to reduction, alkylation, and proteolytic digestion as previously described. All reactions were quenched by adding 1 µL of formic acid and diluted to a volume of 200 µL prior to desalting. Addition of formic acid also caused sodium deoxycholate to precipitate out of solution, and the resulting supernatant was transferred to a new tube before desalting. Desalting and sample injection into the mass spectrometer was performed as previously described in the GlycoFASP methods section.

### Unenriched tear fluid processing

Proteins extracted from tear fluid were diluted to a final concentration of 0.2 mg/mL in 100 µL of PBS. DTT was then added to a concentration of 2 mM and reacted at 65 °C for 1 hour followed by alkylation in 5 mM IAA for 15 min in the dark at RT. Subsequently, digestion with trypsin was done at a 1:50 enzyme:substrate ratio for 6 hours at 37 °C. All reactions were quenched by adding 1 µL of formic acid and diluted to a volume of 200 µL prior to desalting. Desalting and sample injection into the mass spectrometer was performed as previously described in the GlycoFASP methods section.

### Mass spectrometry data acquisition

Samples were analyzed by online nanoflow liquid chromatography-tandem mass spectrometry using an Orbitrap Eclipse Tribrid mass spectrometer (Thermo Fisher Scientific) coupled to a Dionex UltiMate 3000 HPLC (Thermo Fisher Scientific). For each analysis, 4 µL was injected onto an Acclaim PepMap 100 column packed with 2 cm of 5 µm C18 material (Thermo Fisher, 164564) using 0.1% formic acid in water (solvent A). Peptides were then separated on a 15 cm PepMap RSLC EASY-Spray C18 column packed with 2 µm C18 material (Thermo Fisher, ES904) using a gradient from 0-35% solvent B (0.1% formic acid with 80% acetonitrile) in 60 min. Full scan MS1 spectra were collected at a resolution of 60,000, an automatic gain control target of 3e5, and a mass range from *m/z* 300 to 1500. Dynamic exclusion was enabled with a repeat count of 2, repeat duration of 7 s, and exclusion duration of 7 s. Only charge states 2 to 6 were selected for fragmentation. MS2s were generated at top speed for 3 seconds. Higher-energy collisional dissociation (HCD) was performed on all selected precursor masses with the following parameters: isolation window of 2 m/z, 29% normalized collision energy, orbitrap detection (resolution of 7,500), maximum inject time of 50 ms, and a standard automatic gain control target. An additional electron transfer dissociation (ETD) fragmentation of the same precursor was triggered if 1) the precursor mass was between m*/z* 300 to 1500 and 2) 3 of 8 HexNAc or NeuAc fingerprint ions (126.055, 138.055, 144.07, 168.065, 186.076, 204.086, 274.092, and 292.103) were present at *m/z* ± 0.1 and greater than 5% relative intensity. Two files were collected for each sample: the first collected an ETD scan with supplemental energy (EThcD) while the second method collected a scan without supplemental energy. Both used charge-calibrated ETD reaction times, 100 ms maximum injection time, and standard injection targets. EThcD parameters were as follows: Orbitrap detection (resolution 7,500), calibrated charge-dependent ETD times, 15% nCE for HCD, maximum inject time of 150 ms, and a standard precursor injection target. For the second file, dependent scans were only triggered for precursors below *m/z* 1000, and data were collected in the ion trap using a normal scan rate.

### Mass spectrometry data analysis

Raw files were searched using Byonic (version 4.5.2, Protein Metrics, Inc.) against the UniProtKB/Swiss-Prot *Homo sapiens* proteome (Query: proteome:up000005640 AND reviewed:true) and a curated mucin database which was generated from a previous study. Briefly, the mucin database consists of nearly 350 proteins from Uniprot’s annotated human proteome predicted to bear the dense O-glycosylation characteristic of mucin domains. For all samples, we used the default O-glycan database containing 9 common structures. Raw files were first searched against the human proteome and then the curated mucin database. In both cases, files were searched with semi-specific cleavage N-terminal to Ser and Thr and six allowed missed cleavages. Samples treated with trypsin were searched with the same parameters but also allowed cleavage C-terminal to Arg or Lys. Mass tolerance was set to 10 ppm for MS1’s and 20 ppm for MS2’s. Met oxidation was set as a variable modification and carbamidomethyl Cys was set as a fixed modification. From the Byonic search results, glycopeptides were filtered to a score of >200 and a logprob of >2. From the remaining list of glycopeptides, the extracted ion chromatograms, full mass spectra (MS1s), and fragmentation spectra (MS2s) were investigated in XCalibur QualBrowser (Thermo) to generate a list of true-positive glycopeptides, as reported **Table 1**. Each reported glycopeptide listed in **Table 1** was manually validated from the filtered list of Byonic’s reported peptides (score>200 and logprob >2) according to the following steps: The MS1 was first used to confirm the precursor mass and chosen isotope was correct. This also allowed us to identify any co-isolated species that could interfere with the MS2s and/or explain unassigned peaks. The HCD and EThcD fragmentation spectra were then investigated to identify sufficient coverage to make a sequence assignment. When possible, multiple MS2 scans were averaged to obtain a stronger spectrum. For HCD, an initial glycopeptide identification was confirmed if the presence of the precursor mass without a glycan present (i.e., Y0), along with coverage of b and y ions without glycosylation. For longer peptides, we required the presence of Y0 and fragments that were expected to be abundant (e.g., N-terminally to Pro, C-terminally to Asp). When the peptide contained a Pro at the C-terminus, the b_n-1_ was considered sufficient. Further, when the sequence contained oxidized Met, the Met loss from the bare mass was considered as representative of the naked peptide mass. We then used electron-based fragmentation MS2 spectra for localization. Here, all plausible localizations were considered, regardless of search result output. We confirmed the presence of fragment ions in ETD or EThcD that were between potential glycosylation sites, if sufficient c/z ions were present then a glycan mass was considered localized. For glycopeptide manual validation, extracted ion chromatograms are evaluated at the MS1 level to determine the charge and m/z of the highest abundance precursor species. Mass spectrometry data files and raw search output can be found on PRIDE with identifier PXD064788.

### Lacritin Protein Modeling with AlphaFold 3.0 and GlycoShape

The most abundant O-glycan structures of lacritin were first calculated at each O-glycosite using LFQ of AUC intensities of XICs of all lacritin glycopeptides identified (**Supplementary data 1**). Once glycan structures were determined, lacritin’s Uniprot ID (Q9GZZ8) was fetched on GlycoShape webserver’s Re-Glyco tab (https://glycoshape.org/reglyco). Corresponding glycans were then manually input using the “Advanced (Site-by-Site) Glycosylation” feature before hitting “process” for a predicted structure. AlphaFold 3.0 was accessed through the AlphaFoldServer website and the canonical sequence (Uniprot ID: Q9GZZ8) of lacritin was input manually. Next, glycan structures were selected using 3-letter Chemical Component Dictionary (CCD) codes. Here, NAG was used for a single HexNAc residue, BGC was used for a beta-linked Hexose residue, and FUC was used for a Fucose residue. The resultant PDB file was loaded into Pymol 2.0 for glycan recoloration and better visualization. For lacritin C-terminal Alpha helix modeling, the GlycoShape PDB file was exported to Pymol 2.0 and only the C-terminal helix (amino acids 108-138) was shown with the rest of the protein sequence marked as hidden. For isoform C-terminus structure analysis, AlphaFold 3.0 was used as described above.

## Supporting information

This article contains supporting information.

## Data availability statement

All mass spectrometry data and search results acquired for this manuscript have been deposited on the PRIDE repository. Reviewers can access raw data with the following login information: Project accession: PXD064788 Token: fO49eEArej9P. Alternatively, reviewer can access the dataset by logging in to the PRIDE website using the following account details: Username: Password: ErCHGVBtOjJw

## Author contributions

V.C, N.G.K., and S.A.M. conceptualization; V.C., I.L., R.C., and N.C. data curation; V.C., I.L., R.C., and N.C. formal analysis; V.C. investigation; V.C., N.G.K., and K.E.M. methodology; V.C. validation; V.C. visualization; V.C. and S.A.M. writing– original draft; V.C., R.C., N.C., N.G.K., and S.A.M. writing–review & editing; N.G.K. and S.A.M. supervision; N.G.K. and S.A.M. funding Acquisition; V.C. and N.G.K. project administration; N.G.K. and S.A.M. resources.

## Funding and additional information

V.C. is supported by an NSF GRFP (DGE-2139841).S.A.M. is supported by CRI Lloyd J Old STAR Award and a NIGMS R35-GM147039. I.L. is supported by The Yale College First-Year Summer Research Fellowship in the Sciences and Engineering. We also gratefully acknowledge financial contribution to NGK from OsloMet’s seeding fund for internationalization 2024 (3000-116) and to TPU from Department of Medical Biochemistry, Oslo University Hospital, Oslo Norway.

## Abbreviations

The abbreviations used are

ACN: Acetonitrile
ETD: Electron transfer dissociation
EThcD: Electron transfer/higher-energy collision dissociation
HCD: Higher-energy collisional dissociation
DED: Dry eye disease
TGM2: Transglutaminase 2
IgA1: ImmunoglobulinA1
IgJ: Immunoglobulin J chain
PBS: Phosphate Buffered Saline
T antigen: Galβ1-3GalNAcα1-Ser/Thr
Tn antigen: GalNAcα1-Ser/Thr. Type 3 H-antigen (Fuc α1-2-Galβ1-3GalNAcα1-Ser/Thr)

## Reference

(1) Green-Church, K. B.; Butovich, I.; Willcox, M.; Borchman, D.; Paulsen, F.; Barabino, S.; Glasgow, B. J. The International Workshop on Meibomian Gland Dysfunction: Report of the Subcommittee on Tear Film Lipids and Lipid-Protein Interactions in Health and Disease. Investig. Ophthalmol. Vis. Sci. 2011, 52 (4), 1979–1993. 10.1167/iovs.10-6997d.

(2) Pflugfelder, S. C.; De Paiva, C. S.; Li, D. Q.; Stern, M. E. Epithelial-Immune Cell Interaction in Dry Eye. Cornea 2008, 27 (SUPPL. 1), 1–7. 10.1097/ICO.0b013e31817f4075.

(3) Pflugfelder, S. C.; Massaro-Giordano, M.; Perez, V. L.; Hamrah, P.; Deng, S. X.; Espandar, L.; Foster, C. S.; Affeldt, J.; Seedor, J. A.; Afshari, N. A.; Chao, W.; Allegretti, M.; Mantelli, F.; Dana, R. Topical Recombinant Human Nerve Growth Factor (Cenegermin) for Neurotrophic Keratopathy: A Multicenter Randomized Vehicle-Controlled Pivotal Trial. Ophthalmology 2020, 127 (1), 14–26. 10.1016/j.ophtha.2019.08.020.

(4) Craig, J. P.; Nichols, K. K.; Akpek, E. K.; Caffery, B.; Dua, H. S.; Joo, C. K.; Liu, Z.; Nelson, J. D.; Nichols, J. J.; Tsubota, K.; Stapleton, F. TFOS DEWS II Definition and Classification Report. Ocular Surface. Elsevier Inc. July 1, 2017, pp 276–283. 10.1016/j.jtos.2017.05.008.

(5) Stapleton, F.; Alves, M.; Bunya, V. Y.; Jalbert, I.; Lekhanont, K.; Malet, F.; Na, K. S.; Schaumberg, D.; Uchino, M.; Vehof, J.; Viso, E.; Vitale, S.; Jones, L. TFOS DEWS II Epidemiology Report. Ocul. Surf. 2017, 15 (3), 334–365. 10.1016/j.jtos.2017.05.003.

(6) Farrand, K. F.; Fridman, M.; Stillman, I. Ö.; Schaumberg, D. A. Prevalence of Diagnosed Dry Eye Disease in the United States Among Adults Aged 18 Years and Older. Am. J. Ophthalmol. 2017, 182, 90–98. 10.1016/j.ajo.2017.06.033.

(7) Azkargorta, M.; Soria, J.; Ojeda, C.; Guzmán, F.; Acera, A.; Iloro, I.; Suárez, T.; Elortza, F. Human Basal Tear Peptidome Characterization by CID, HCD, and ETD Followed by in Silico and in Vitro Analyses for Antimicrobial Peptide Identification. J. Proteome Res. 2015, 14 (6), 2649–2658. 10.1021/acs.jproteome.5b00179.

(8) Ali, N.; Turkiewicz, A.; Hughes, V.; Folkesson, E.; Tjörnstand, J.; Neuman, P.; Önnerfjord, P.; Englund, M. Proteomics Profiling of Human Synovial Fluid Suggests Increased Protein Interplay in Early-Osteoarthritis (OA) That Is Lost in Late-Stage OA. Mol. Cell. Proteomics 2022, 21 (3). 10.1016/j.mcpro.2022.100200.

(9) Samudre, S.; Lattanzio, F. A.; Lossen, V.; Hosseini, A.; Sheppard, J. D.; McKown, R. L.; Laurie, G. W.; Williams, P. B. Lacritin, a Novel Human Tear Glycoprotein, Promotes Sustained Basal Tearing and Is Well Tolerated. Investig. Ophthalmol. Vis. Sci. 2011, 52 (9), 6265–6270. 10.1167/iovs.10-6220.

(10) Conley, S. M.; Naash, M. I. Focus on Molecules: Lacritin. Exp. Eye Res. 2009, 89 (3), 278–279. 10.1016/j.exer.2009.03.023.

(11) Samudre, S.; Lattanzio, F. A.; Lossen, V.; Hosseini, A.; Sheppard, J. D.; McKown, R. L.; Laurie, G. W.; Williams, P. B. Lacritin, a Novel Human Tear Glycoprotein, Promotes Sustained Basal Tearing and Is Well Tolerated. Investig. Ophthalmol. Vis. Sci. 2011, 52 (9), 6265–6270. 10.1167/iovs.10-6220.

(12) Karnati, R.; Laurie, D. E.; Laurie, G. W. Lacritin and the Tear Proteome as Natural Replacement Therapy for Dry Eye. Exp. Eye Res. 2013, 117, 39–52. 10.1016/j.exer.2013.05.020.

(13) Efraim, Y.; Chen, F. Y. T.; Cheong, K. N.; Gaylord, E. A.; McNamara, N. A.; Knox, S. M. A Synthetic Tear Protein Resolves Dry Eye through Promoting Corneal Nerve Regeneration. Cell Rep. 2022, 40 (9). 10.1016/j.celrep.2022.111307.

(14) Tauber, J.; Laurie, G. W.; Parsons, E. C.; Odrich, M. G.; Abrams, M. A.; Asbell, P. A.; Berdy, G. J.; Bergstrom, L. K.; Bowden, F. W.; Bower, K. S.; Boyce, J. D.; Davanzo, R. J.; El-Harazi, L.; Epitropoulos, A. T.; Forstot, S. L.; Goldberg, D. F.; Greiner, J. V.; Haider, K. M.; Hardten, D. R.; Hauswirth, S. G.; Hom, M. M.; Kelley, K. A.; Khachikian, S. S.; Kim, J. L.; Majmudar, P. A.; Martel, J. R.; Massaro-Giordano, G.; McGehee, T. W.; McNamara, N. A.; Meyer, J. C.; Nichols, K. K.; Perez, B. R.; Rubin, M. S.; Sall, K.; Schultze, R. L.; Schwartz, J. L.; Segal, B.; Sheppard, J. D.; Tauber, J.; Thompson, V. M.; Tims, J. S.; Van, D. T. Lacripep for the Treatment of Primary Sjögren-Associated Ocular Surface Disease: Results of the First-In-Human Study. Cornea 2023, 42 (7), 847–857. 10.1097/ICO.0000000000003091.

(15) Dias-Teixeira, K.; Horton, X.; McKown, R.; Romano, J.; Laurie, G. W. The Lacritin-Syndecan-1-Heparanase Axis in Dry Eye Disease. In Advances in Experimental Medicine and Biology; Springer, 2020; Vol. 1221, pp 747–757. 10.1007/978-3-030-34521-1_31.

(16) Ma, P.; Beck, S. L.; Raab, R. W.; McKown, R. L.; Coffman, G. L.; Utani, A.; Chirico, W. J.; Rapraeger, A. C.; Laurie, G. W. Heparanase Deglycanation of Syndecan-1 Is Required for Binding of the Epithelial-Restricted Prosecretory Mitogen Lacritin. J. Cell Biol. 2006, 174 (7), 1097–1106. 10.1083/jcb.200511134.

(17) Francisco, V. V.; Romano, J. A.; McKown, R. L.; Green, K.; Zhang, L.; Raab, R. W.; Ryan, D. S.; Hutnik, C. M. L.; Frierson, H. F.; Laurie, G. W. Tissue Transglutaminase Is a Negative Regulator of Monomeric Lacritin Bioactivity. Investig. Ophthalmol. Vis. Sci. 2013, 54 (3), 2123–2132. 10.1167/iovs.12-11488.

(18) Aragona, P.; Aguennouz, M.; Rania, L.; Postorino, E.; Sommario, M. S.; Roszkowska, A. M.; De Pasquale, M. G.; Pisani, A.; Puzzolo, D. Matrix Metalloproteinase 9 and Transglutaminase 2 Expression at the Ocular Surface in Patients with Different Forms of Dry Eye Disease. Ophthalmology 2015, 122 (1), 62–71. 10.1016/j.ophtha.2014.07.048.

(19) Tovar-Vidales, T.; Roque, R.; Clark, A. F.; Wordinger, R. J. Tissue Transglutaminase Expression and Activity in Normal and Glaucomatous Human Trabecular Meshwork Cells and Tissues. Investig. Ophthalmol. Vis. Sci. 2008, 49 (2), 622–628. 10.1167/iovs.07-0835.

(20) Mckown, R. L.; Wang, N.; Raab, R. W.; Karnati, R.; Zhang, Y.; Williams, P. B.; Laurie, G. W. Lacritin and Other New Proteins of the Lacrimal Functional Unit. 2010, 88 (5), 848–858. 10.1016/j.exer.2008.09.002.Lacritin.

(21) Movassaghi, C. S.; Sun, J.; Jiang, Y.; Turner, N.; Chang, V.; Chung, N.; Chen, R. J.; Browne, E. N.; Lin, C.; Schweppe, D. K.; Malaker, S. A.; Meyer, J. G. Recent Advances in Mass Spectrometry-Based Bottom-Up Proteomics. Anal. Chem. 2025. 10.1021/acs.analchem.4c06750.

(22) Bagdonaite, I.; Malaker, S. A.; Polasky, D. A.; Riley, N. M.; Schjoldager, K.; Vakhrushev, S. Y.; Halim, A.; Aoki-Kinoshita, K. F.; Nesvizhskii, A. I.; Bertozzi, C. R.; Wandall, H. H.; Parker, B. L.; Thaysen-Andersen, M.; Scott, N. E. Glycoproteomics. Nat. Rev. Methods Prim. 2022, 2 (1). 10.1038/s43586-022-00128-4.

(23) Nguyen-Khuong, T.; Everest-Dass, A. V.; Kautto, L.; Zhao, Z.; Willcox, M. D. P.; Packer, N. H. Glycomic Characterization of Basal Tears and Changes with Diabetes and Diabetic Retinopathy. Glycobiology 2013, 25 (3), 269–283. 10.1093/glycob/cwu108.

(24) Messina, A.; Palmigiano, A.; Tosto, C.; Romeo, D. A.; Sturiale, L.; Garozzo, D.; Leonardi, A. Tear N-Glycomics in Vernal and Atopic Keratoconjunctivit. *Allergy Eur*. J. Allergy Clin. Immunol. 2021, 76 (8), 2500–2509. 10.1111/all.14775.

(25) Zhou, L.; Beuerman, R. W.; Chew, A. P.; Koh, S. K.; Cafaro, T. A.; Urrets-Zavalia, E. A.; Urrets-Zavalia, J. A.; Li, S. F. Y.; Serra, H. M. Quantitative Analysis of N-Linked Glycoproteins in Tear Fluid of Climatic Droplet Keratopathy by Glycopeptide Capture and ITRAQ. J. Proteome Res. 2009, 8 (4), 1992–2003. 10.1021/pr800962q.

(26) Schmelter, C.; Brueck, A.; Perumal, N.; Qu, S.; Pfeiffer, N.; Grus, F. H. Lectin-Based Affinity Enrichment and Characterization of N-Glycoproteins from Human Tear Film by Mass Spectrometry. Molecules 2023, 28 (2). 10.3390/molecules28020648.

(27) Malaker, S. A.; Riley, N. M.; Shon, D. J.; Pedram, K.; Krishnan, V.; Dorigo, O.; Bertozzi, C. R. Revealing the Human Mucinome. Nat. Commun. 2022, 13 (1). 10.1038/s41467-022-31062-4.

(28) Shon, D. J.; Malaker, S. A.; Pedram, K.; Yang, E.; Krishnan, V.; Dorigo, O.; Bertozzi, C. R. An Enzymatic Toolkit for Selective Proteolysis, Detection, and Visualization of Mucin-Domain Glycoproteins. Proc. Natl. Acad. Sci. U. S. A. 2020, 117 (35), 21299–21307. 10.1073/pnas.2012196117.

(29) Chongsaritsinsuk, J.; Steigmeyer, A. D.; Mahoney, K. E.; Rosenfeld, M. A.; Lucas, T. M.; Smith, C. M.; Li, A.; Ince, D.; Kearns, F. L.; Battison, A. S.; Hollenhorst, M. A.; Judy Shon, D.; Tiemeyer, K. H.; Attah, V.; Kwon, C.; Bertozzi, C. R.; Ferracane, M. J.; Lemmon, M. A.; Amaro, R. E.; Malaker, S. A. Glycoproteomic Landscape and Structural Dynamics of TIM Family Immune Checkpoints Enabled by Mucinase SmE. Nat. Commun. 2023, 14 (1). 10.1038/s41467-023-41756-y.

(30) Mahoney, K. E.; Chang, V.; Lucas, T. M.; Maruszko, K.; Malaker, S. A. Mass Spectrometry-Compatible Elution Technique Enables an Improved Mucin-Selective Enrichment Strategy to Probe the Mucinome. Anal. Chem. 2023. 10.1021/acs.analchem.3c05762.

(31) Shane M. Finn; Keira E. Mahoney; Taryn M. Lucas; Valentina Rangel-Angarita; Ryan J. Chen; Stacy A. Malaker. GlycoFASP: A Universal Method to Prepare Complex Mixtures for O-Glycoproteomic Analysis. ChemRxiv 2025. 10.7868/80424857017030112.

(32) Mahoney, K. E.; Chang, V.; Lucas, T. M.; Maruszko, K.; Malaker, S. A. Mass Spectrometry-Compatible Elution Technique Enables an Improved Mucin-Selective Enrichment Strategy to Probe the Mucinome. Anal. Chem. 2024, 96 (13), 5242–5250. 10.1021/acs.analchem.3c05762.

(33) Tham, M. L.; Mahmud, A.; Abdullah, M.; Md Saleh, R.; Mohammad Razali, A.; Cheah, Y. K.; Mohd Taib, N.; Ho, K. L.; Mahmud, M.; Mohd Isa, M. Tear Samples for Protein Extraction: Comparative Analysis of Schirmer’s Test Strip and Microcapillary Tube Methods. Cureus 2023, 15 (12). 10.7759/cureus.50972.

(34) Mantelli, F.; Argüeso, P. Functions of Ocular Surface Mucins in Health and Disease. Curr. Opin. Allergy Clin. Immunol. 2008, 8 (5), 477–483. 10.1097/ACI.0b013e32830e6b04.

(35) Nguyen-Khuong, T.; Everest-Dass, A. V.; Kautto, L.; Zhao, Z.; Willcox, M. D. P.; Packer, N. H. Glycomic Characterization of Basal Tears and Changes with Diabetes and Diabetic Retinopathy. Glycobiology 2013, 25 (3), 269–283. 10.1093/glycob/cwu108.

(36) Schulz, B. L.; Oxley, D.; Packer, N. H.; Karlsson, N. G. Identification of Two Highly Sialylated Human Tear-Fluid DMBT1 Isoforms : The Major High-Molecular-Mass Glycoproteins in Human Tears; 2002; Vol. 366.

(37) Azuma, M.; Application, F.; Data, P. Partial Peptide of Lacritin. 2013, 2 (12).

(38) Morimoto-Tochigi, A.; Nakajima, T.; Fujii, A.; Shearer, T. R.; Azuma, M. Functions of Primate Lacritin: Enhanced Secretion of Tear Proteins From Lacrimal Acinar Cells and Promotion of Corneal Epithelial Cell Adhesion. Invest. Ophthalmol. Vis. Sci. 2010, 51 (13), 4178.

(39) Zhang, Y.; Wang, N.; Raab, R. W.; McKown, R. L.; Irwin, J. A.; Kwon, I.; Van Kuppevelt, T. H.; Laurie, G. W. Targeting of Heparanase-Modified Syndecan-1 by Prosecretory Mitogen Lacritin Requires Conserved Core GAGAL plus Heparan and Chondroitin Sulfate as a Novel Hybrid Binding Site That Enhances Selectivity. J. Biol. Chem. 2013, 288 (17), 12090–12101. 10.1074/jbc.M112.422717.

(40) Tropea, B.; Fadda, E.; Ives, C. M.; Singh, O.; D’Andrea, S.; Fogarty, C. A.; Harbison, A. M.; Satheesan, A.; Tropea, B.; Fadda, E. Restoring Protein Glycosylation with GlycoShape. Nat. Methods 2024, 21 (November), 2023.12.11.571101. 10.1038/s41592-024-02464-7.

(41) Ince, D.; Lucas, T. M.; Malaker, S. A. Current Strategies for Characterization of Mucin-Domain Glycoproteins. Current Opinion in Chemical Biology. Elsevier Ltd August 1, 2022. 10.1016/j.cbpa.2022.102174.

(42) Kearns, F. L.; Rosenfeld, M. A.; Amaro, R. E. Breaking Down the Bottlebrush: Atomically-Detailed Structural Dynamics of Mucins. bioRxiv 2024, 2024.04.09.588790. 10.1021/acs.jcim.4c00613.

(43) Georgiev, G. A.; Gh, M. S.; Romano, J.; Teixeira, K. L. D.; Struble, C.; Ryan, D. S.; Sia, R. K.; Kitt, J. P.; Harris, J. M.; Hsu, K. L.; Libby, A.; Odrich, M. G.; Suárez, T.; McKown, R. L.; Laurie, G. W. Lacritin Proteoforms Prevent Tear Film Collapse and Maintain Epithelial Homeostasis. J. Biol. Chem. 2021, 296, 1–16. 10.1074/jbc.RA120.015833.

(44) Lee, C.; Kim, M.; Park, C.; Jo, W.; Seo, J. K.; Kim, S.; Oh, J.; Kim, C. S.; Ryu, H. S.; Lee, K. H.; Park, J. Epigenetic Regulation of Neuregulin 1 Promotes Breast Cancer Progression Associated to Hyperglycemia. Nat. Commun. 2023, 14 (1). 10.1038/s41467-023-36179-8.

(45) Vullhorst, D.; Ahmad, T.; Karavanova, I.; Keating, C.; Buonanno, A. Structural Similarities between Neuregulin 1–3 Isoforms Determine Their Subcellular Distribution and Signaling Mode in Central Neurons. J. Neurosci. 2017, 37 (21), 5232–5249. 10.1523/JNEUROSCI.2630-16.2017.

(46) Fleck, D.; Voss, M.; Brankatschk, B.; Giudici, C.; Hampel, H.; Schwenk, B.; Edbauer, D.; Fukumori, A.; Steiner, H.; Kremmer, E.; Haug-Kröper, M.; Rossner, M. J.; Fluhrer, R.; Willem, M.; Haass, C. Proteolytic Processing of Neuregulin 1 Type III by Three Intramembrane-Cleaving Proteases. J. Biol. Chem. 2016, 291 (1), 318–333. 10.1074/jbc.M115.697995.

(47) Afshari, A. R.; Chang, V.; Thomsson, K. A.; Höglund, J.; Browne, E. N.; Karadzhov, G.; Mahoney, K. E.; Lucas, T. M.; Rangel-Angarita, V.; Ryberg, H.; Gidwani, K.; Pettersson, K.; Rolfson, O.; Björkman, L. I.; Eisler, T.; Schmidt, T. A.; Jay, G. D.; Malaker, S. A.; Karlsson, N. G. Glycoproteoforms of Osteoarthritis-Associated Lubricin in Plasma and Synovial Fluid. Mol. Cell. Proteomics 2025, 24 (3), 100923. 10.1016/j.mcpro.2025.100923.

(48) Justis, B. M.; Coburn, C. E.; Tyler, E. M.; Showalter, R. S.; Dissler, B. J.; Li, M.; McNamara, N. A.; Laurie, G. W.; McKown, R. L. Development of a Quantitative Immunoassay for Tear Lacritin Proteoforms. Transl. Vis. Sci. Technol. 2020, 9 (9), 1–12. 10.1167/tvst.9.9.13.

(49) Goth, C. K.; Halim, A.; Khetarpal, S. A.; Rader, D. J.; Clausen, H.; Schjoldager, K. T. B. G. A Systematic Study of Modulation of ADAM-Mediated Ectodomain Shedding by Site-Specific O-Glycosylation. Proc. Natl. Acad. Sci. U. S. A. 2015, 112 (47), 14623–14628. 10.1073/pnas.1511175112.

(50) Riethmueller, S.; Somasundaram, P.; Ehlers, J. C.; Hung, C. W.; Flynn, C. M.; Lokau, J.; Agthe, M.; Düsterhöft, S.; Zhu, Y.; Grötzinger, J.; Lorenzen, I.; Koudelka, T.; Yamamoto, K.; Pickhinke, U.; Wichert, R.; Becker-Pauly, C.; Rädisch, M.; Albrecht, A.; Hessefort, M.; Stahnke, D.; Unverzagt, C.; Rose-John, S.; Tholey, A.; Garbers, C. Proteolytic Origin of the Soluble Human IL-6R In Vivo and a Decisive Role of N-Glycosylation; 2017; Vol. 15. 10.1371/journal.pbio.2000080.

(51) Hrdličková-Cela, E.; Plzák, J.; Smetana, K.; Mělková, Z.; Kaltner, H.; Filipec, M.; Liu, F.-T.; Gabius, H.-J. Detection of Galectin-3 in Tear Fluid at Disease States and Immunohistochemical and Lectin Histochemical Analysis in Human Corneal and Conjunctival Epithelium. Br. J. Ophthalmol. 2001, 85 (11), 1336 LP – 1340. 10.1136/bjo.85.11.1336.

(52) Guzman-Aranguez, A.; Mantelli, F.; Argüeso, P. Mucin-Type O-Glycans in Tears of Normal Subjects and Patients with Non-Sjögren’s Dry Eye. Investig. Ophthalmol. Vis. Sci. 2009, 50 (10), 4581–4587. 10.1167/iovs.09-3563.

(53) An, H. J.; Ninonuevo, M.; Aguilan, J.; Liu, H.; Lebrilla, C. B.; Alvarenga, L. S.; Mannis, M. J. Glycomics Analyses of Tear Fluid for the Diagnostic Detection of Ocular Rosacea. J. Proteome Res. 2005, 4 (6), 1981–1987. 10.1021/pr0501620.

(54) Ballard, C. J.; Smutny, M. R.; Chau, L. D.; Wong, C. K.; Aharoni, H. M.; Lee, H. K.; Chapla, D. G.; Hurtado-Guerrero, R.; Moremen, K. W.; Gerken, T. A. Charge Matters: How Flanking Substrate Charge Modulates O-Glycan Core Elongation. Glycobiology 2025, 35 (5). 10.1093/glycob/cwaf014.

(55) Zhou, L.; Zhao, S. Z.; Koh, S. K.; Chen, L.; Vaz, C.; Tanavde, V.; Li, X. R.; Beuerman, R. W. In-Depth Analysis of the Human Tear Proteome. J. Proteomics 2012, 75 (13), 3877–3885. 10.1016/j.jprot.2012.04.053.

(56) Jung, J. H.; Ji, Y. W.; Hwang, H. S.; Oh, J. W.; Kim, H. C.; Lee, H. K.; Kim, K. P. Proteomic Analysis of Human Lacrimal and Tear Fluid in Dry Eye Disease. Sci. Rep. 2017, 7 (1). 10.1038/s41598-017-13817-y.

(57) Iwasaki, H.; Zhang, Y.; Tachibana, K.; Gotoh, M.; Kikuchi, N.; Kwon, Y. D.; Togayachi, A.; Kudo, T.; Kubota, T.; Narimatsu, H. Initiation of O-Glycan Synthesis in IgA1 Hinge Region Is Determined by a Single Enzyme, UDP-N-Acetyl-α-D-Galactosamine: Polypeptide N-Acetylgalactosaminyltransferase 2. J. Biol. Chem. 2003, 278 (8), 5613–5621. 10.1074/jbc.M211097200.

(58) Endo, T.; Mestecky, J.; Kulhavy, R.; Kobata, A. Carbohydrate Heterogeneity of Human Myeloma Proteins of the IgA1 and IgA2 Subclasses. Mol. Immunol. 1994, 31 (18), 1415–1422. 10.1016/0161-5890(94)90157-0.

(59) Ding, L.; Chen, X.; Cheng, H.; Zhang, T.; Li, Z. Advances in IgA Glycosylation and Its Correlation with Diseases. Front. Chem. 2022, 10 (September), 1–17. 10.3389/fchem.2022.974854.

(60) Dotz, V.; Visconti, A.; Lomax-Browne, H. J.; Clerc, F.; Hipgrave Ederveen, A. L.; Medjeral-Thomas, N. R.; Cook, H. T.; Pickering, M. C.; Wuhrer, M.; Falchi, M. O- And N-Glycosylation of Serum Immunoglobulin a Is Associated with Iga Nephropathy and Glomerular Function. J. Am. Soc. Nephrol. 2021, 32 (10), 2455–2465. 10.1681/ASN.2020081208.

(61) Suzuki, H.; Yasutake, J.; Makita, Y.; Tanbo, Y.; Yamasaki, K.; Sofue, T.; Kano, T.; Suzuki, Y. IgA Nephropathy and IgA Vasculitis with Nephritis Have a Shared Feature Involving Galactose-Deficient IgA1-Oriented Pathogenesis. Kidney Int. 2018, 93 (3), 700–705. 10.1016/j.kint.2017.10.019.

(62) Suzuki, T.; Kawaguchi, A.; Ainai, A.; Tamura, S. I.; Ito, R.; Multihartina, P.; Setiawaty, V.; Pangesti, K. N. A.; Odagiri, T.; Tashiro, M.; Hasegawa, H. Relationship of the Quaternary Structure of Human Secretory IgA to Neutralization of Influenza Virus. Proc. Natl. Acad. Sci. U. S. A. 2015, 112 (25), 7809–7814. 10.1073/pnas.1503885112.

(63) Lomax-Browne, H. J.; Robertson, C.; Antonopoulos, A.; Leathem, A. J. C.; Haslam, S. M.; Dell, A.; Dwek, M. V. Serum IgA1 Shows Increased Levels of A2,6-Linked Sialic Acid in Breast Cancer. Interface Focus 2019, 9 (2). 10.1098/rsfs.2018.0079.

(64) Zhang, S.; Cao, X.; Liu, C.; Li, W.; Zeng, W.; Li, B.; Chi, H.; Liu, M.; Qin, X.; Tang, L.; Yan, G.; Ge, Z.; Liu, Y.; Gao, Q.; Lu, H. N-Glycopeptide Signatures of IgA2 in Serum from Patients with Hepatitis b Virus-Related Liver Diseases. Mol. Cell. Proteomics 2019, 18 (11), 2262–2272. 10.1074/mcp.RA119.001722.

(65) Tarelli, E.; Smith, A. C.; Hendry, B. M.; Challacombe, S. J.; Pouria, S. Human Serum IgA1 Is Substituted with up to Six O-Glycans as Shown by Matrix Assisted Laser Desorption Ionisation Time-of-Flight Mass Spectrometry. Carbohydr. Res. 2004, 339 (13), 2329–2335. 10.1016/j.carres.2004.07.011.

(66) Takahashi, K.; Wall, S. B.; Suzuki, H.; Smith IV, A. D.; Hall, S.; Poulsen, K.; Kilian, M.; Mobley, J. A.; Julian, B. A.; Mestecky, J.; Novak, J.; Renfrow, M. B. Clustered O-Glycans of IgA1: Defining Macro- and Microheterogeneity by Use of Electron Capture/Transfer Dissociation. Mol. Cell. Proteomics 2010, 9 (11), 2545–2557. 10.1074/mcp.M110.001834.

(67) Franc, V.; Řehulka, P.; Raus, M.; Stulík, J.; Novak, J.; Renfrow, M. B.; Šebela, M. Elucidating Heterogeneity of IgA1 Hinge-Region O-Glycosylation by Use of MALDI-TOF/TOF Mass Spectrometry: Role of Cysteine Alkylation during Sample Processing. J. Proteomics 2013, 92, 299–312. 10.1016/j.jprot.2013.07.013.

(68) Gomes, M. M.; Wall, S. B.; Takahashi, K.; Novak, J.; Renfrow, M. B.; Herr, A. B. Analysis of IgA1 N-Glycosylation and Its Contribution to FcαRI Binding. Biochemistry 2008, 47 (43), 11285–11299. 10.1021/bi801185b.

(69) Plomp, R.; De Haan, N.; Bondt, A.; Murli, J.; Dotz, V.; Wuhrer, M. Comparative Glycomics of Immunoglobulin A and G from Saliva and Plasma Reveals Biomarker Potential. Front. Immunol. 2018, 9 (OCT), 1–12. 10.3389/fimmu.2018.02436.

(70) Karav, S.; German, J. B.; Rouquié, C.; Le Parc, A.; Barile, D. Studying Lactoferrin N-Glycosylation. Int. J. Mol. Sci. 2017, 18 (4), 1–14. 10.3390/ijms18040870.

(71) Flanagan, J. L.; Willcox, M. D. P. Role of Lactoferrin in the Tear Film. Biochimie 2009, 91 (1), 35–43. 10.1016/j.biochi.2008.07.007.

(72) Tsai, C. Y.; Hong, C.; Hsu, M. Y.; Lai, T. T.; Huang, C. W.; Lu, C. Y.; Chen, W. L.; Cheng, C. M. Fluorescence-Based Reagent and Spectrum-Based Optical Reader for Lactoferrin Detection in Tears: Differentiating Sjögren’s Syndrome from Non-Sjögren’s Dry Eye Syndrome. Sci. Rep. 2024, 14 (1), 1–8. 10.1038/s41598-024-65487-2.

