## Supplemental Information for "In-depth analysis of the tear fluid glycoproteome reveals diverse lacritin glycosylation and spliceoforms"

**Supplementary Table**

Supplementary Table 1: N-/O-glycoproteomic analysis of tear fluid with GlycoFASP

**Supplementary Figures**

Supplementary Figure 1. GlycoShape modeling of Lacritin C-terminal helix with and without N-glycosylation
Supplementary Figure 2. Lacritin N-glycosylation analysis
Supplementary Figure 3. Lacritin isoform A-specific peptide quantification
Supplementary Figure 4. AlphaFold 3.0 modeling of alpha helix in Lacritin isoforms A,C, and D.
Supplementary Figure 5. Evolutionary analysis of Lacritin isoforms and mucin domain
Supplementary Figure 6. Lactotransferrin N-glycosylation analysis

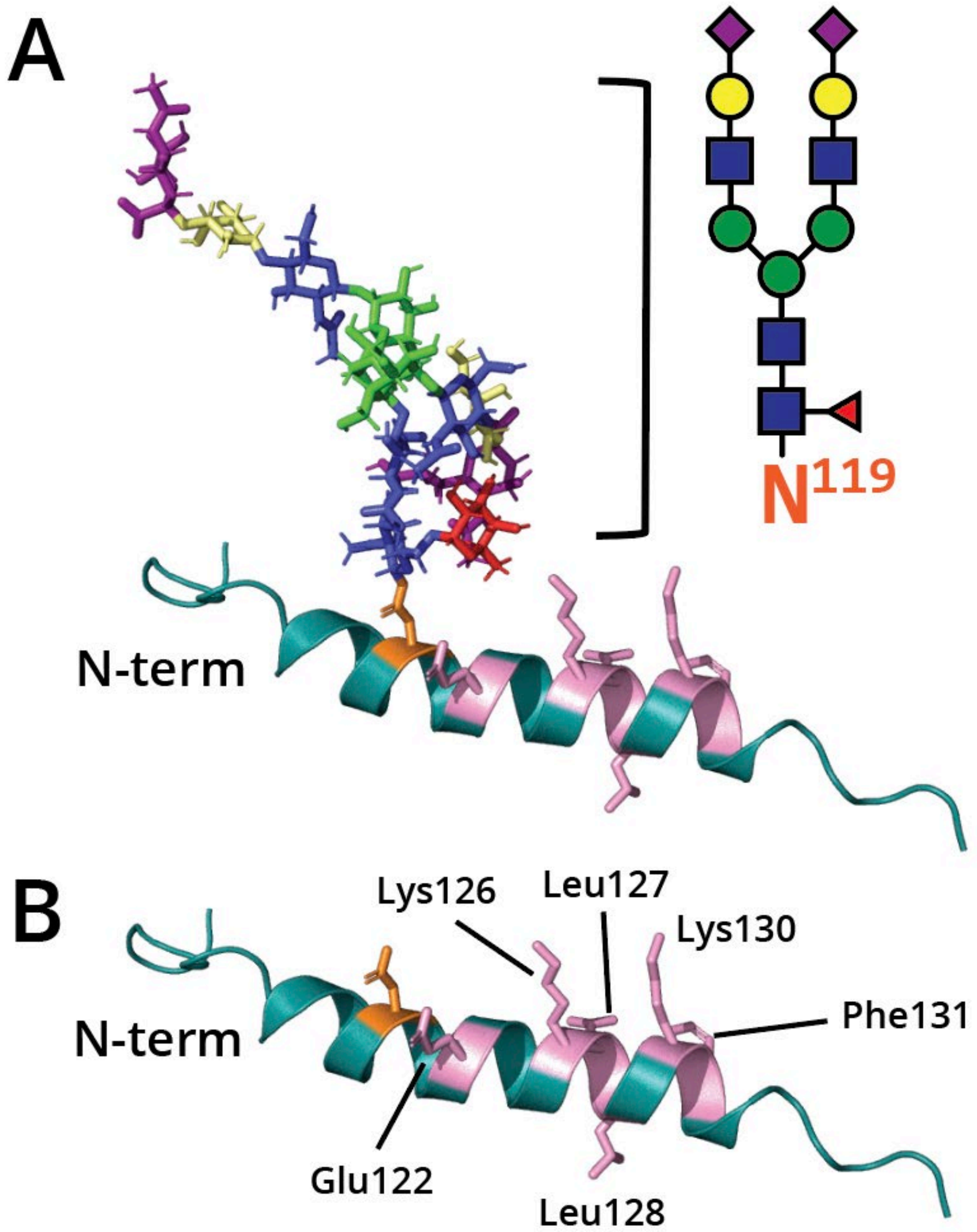


**Supplementary Figure 1. GlycoShape modeling of Lacritin C-terminal helix with and without N-glycosylation at Asp119**.

Amino acid residues 108-138 of Lacritin modeled using GlycoShape. (**A**) C-terminal alpha helix containing an H5N4A2F1 N-glycan on Asp119 (orange). (**B**) C-terminal alpha helix without N-glycan on Asp 119 (orange). Glu122, Lys126, Leu127, Leu128, Lys130, and Phe131 (colored in pink) are previously reported residues which are necessary for syndecan-1 binding.

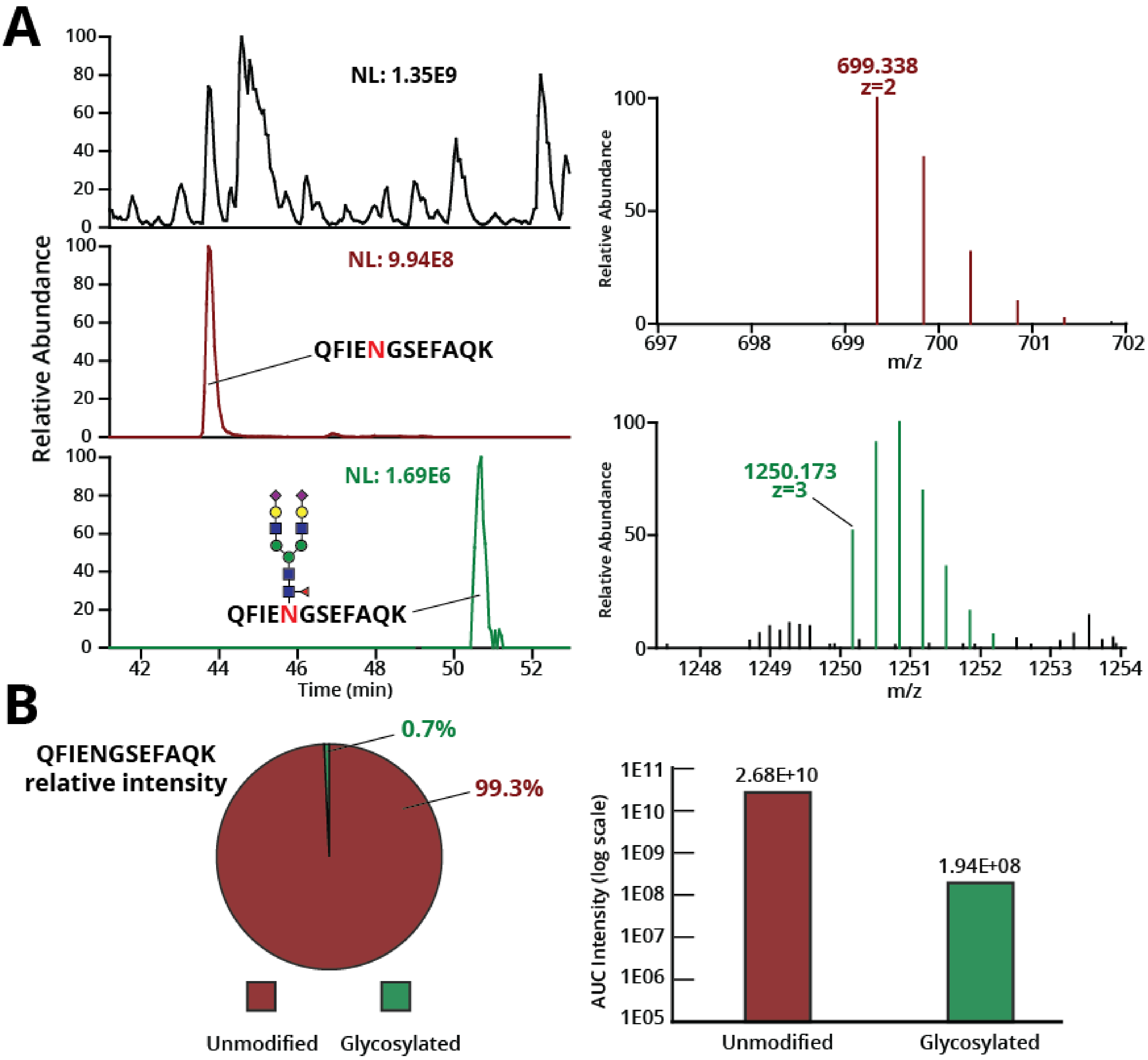


**Supplementary Figure 2. Lacritin N-glycosylation analysis.**
(**A**) Base peak chromarogram (top, left) and XICs of the unmodified peptide (middle, left) bearing lacritin N-glycosite (m/z 699.338, z=2, retention time 42-52 minutes) and the glycosylated peptide(middle, bottom) (m/z 1250.173, z=3, retention time 42-52 minutes). The MS1 of the monoisotopic precursor and its isotopes are shown (right) along with other co-isolated species. (**B**) Quantification of N-glycosite relative occupancy based on LFQ of AUC intensities of XICs.

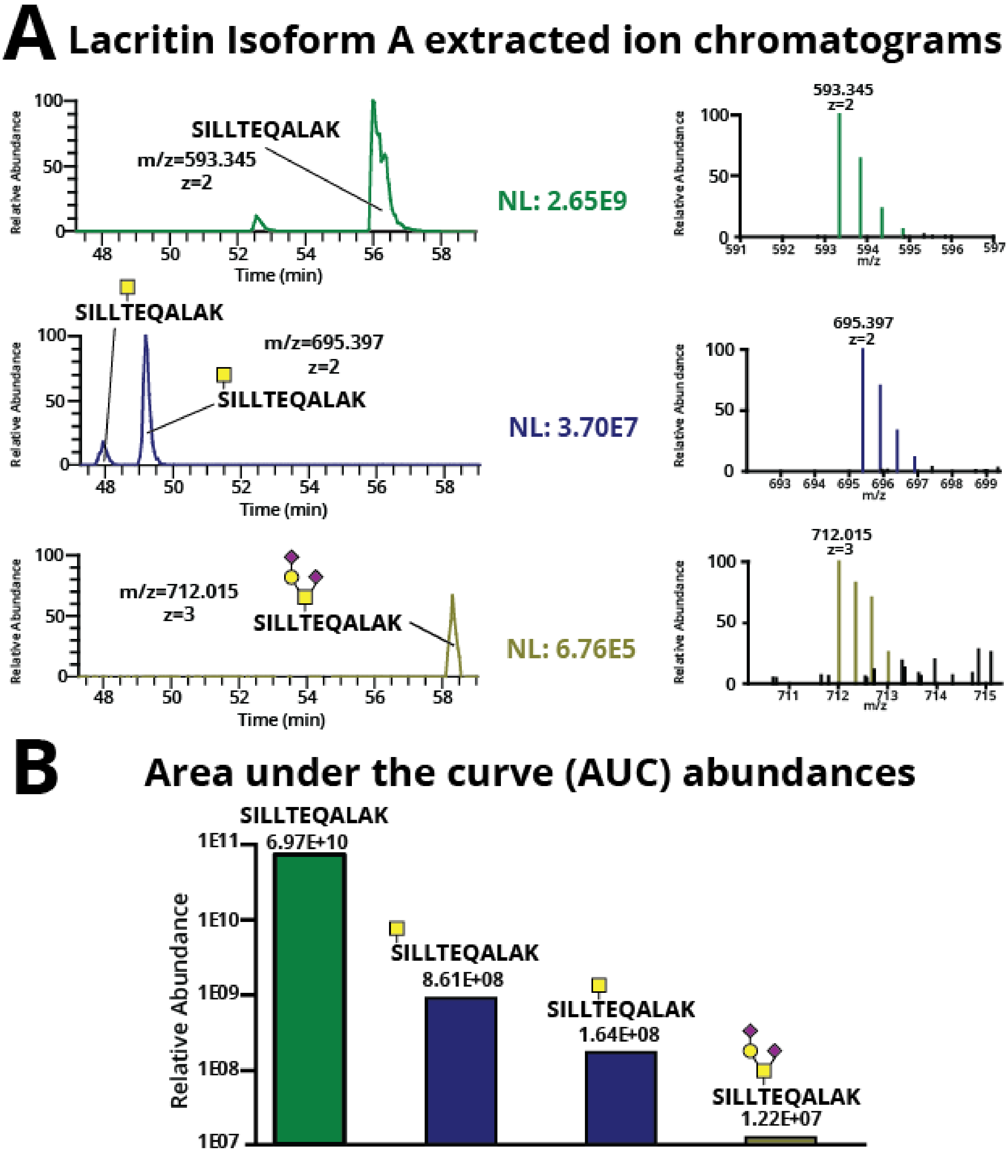


**Supplementary Figure 3. Lacritin isoform A-specific peptide quantification**

(**A**) Lacritin isoform A-specific peptide XICs of the unmodified glycoform (top, m/z 593.345, z=2, retention time 42-52 minutes) and the modified glycoforms (middle, bottom) (m/z 695.397, z=2, and m/z 12.015, z=3, retention time 42-52 minutes). (**B**) Quantification of O-glycosite relative glycosylation frequency based on LFQ of AUC intensities of XICs.

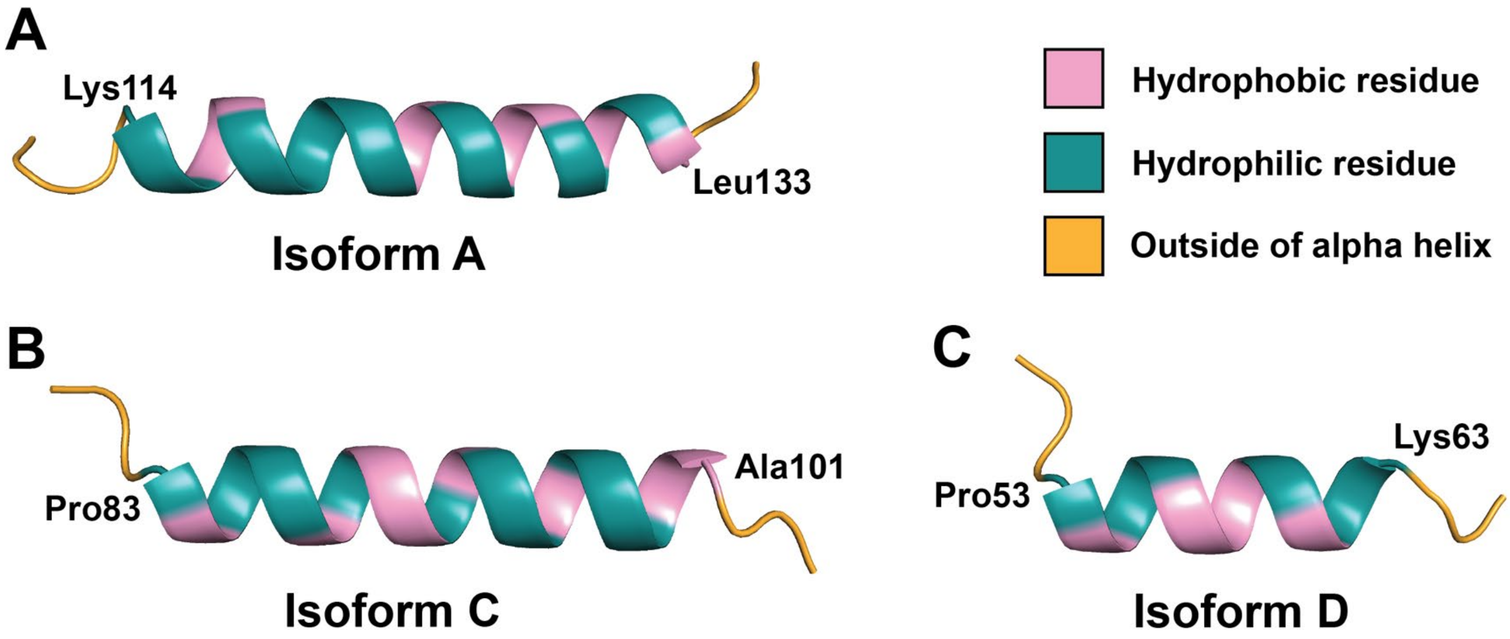


**Supplementary Figure 4. AlphaFold 3.0 modeling of alpha helix in Lacritin isoforms A,C, and D**. In each model, hydrophilic residues are indicated in light blue, hydrophobic residues are indicated in orange, and residues outside the helix are indicated in pink. The total hydropathy score for each alpha helix was calculated using the Kyte-Doolittle scale, where a score >0 indicates hydrophobic and a score <0 indicates hydrophilic. The total score represents the sum of the Kyte-Doolittle values for all residues within the helix. (A) AlphaFold predicted structure of Isoform A alpha helix spanning Lys114 to Leu133.The total hydropathy score is -9.0, indicating that this helix is the most hydrophilic among the helices from three above isoforms. (B) AlphaFold predicted structure of Isoform C alpha helix spanning Pro83 to Ala101. The total hydropathy score is 1.2, suggesting a overall hydrophobic helix. (C) AlphaFold predicted structure of Isoform D alpha helix spanning Pro53 to Lys63. The total hydropathy score is -4.2, indicating a predominantly hydrophilic segment.

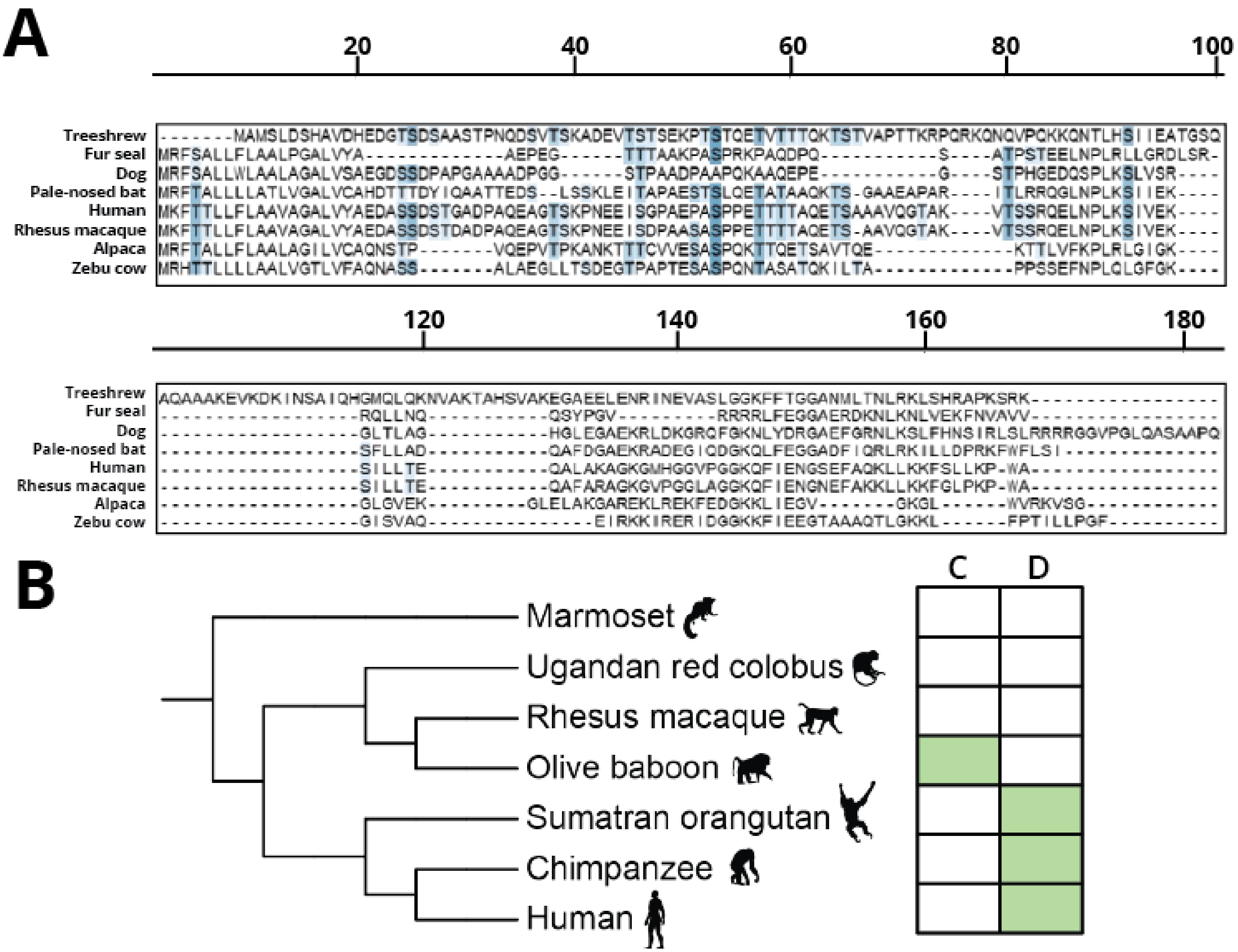


**Supplementary Figure 5. Multiple sequence alignment of lacritin orthologs shows N-terminal mucin domain O-glycosite conservation and differential isoform conservation in primates.**
**(A)** Orthologous sequences of lacritin were collected from the NCBI Ortholog database (LACRT) and InParanoidDB9 (LACRT) to sample phylogenetic diversity among mammalian species. Sequences were obtained for the following species with RefSeq or UniProt IDs indicated: Treeshrew (*Tupaia chinensis*, XP_006169178.1), Northern Fur Seal (*Callorhinus ursinus*, A0A3Q7PZI8), Dog (*Canis lupus familiaris*, XP_038370182.1), Pale Spear-nosed Bat (*Phyllostomus discolor*, A0A7E6D6B7), Human (*Homo sapiens*, Q9GZZ8), Rhesus Macaque (*Macaca mulatta*, Q2Q5T4), Alpaca (*Vicugna pacos*, A0A6J3AZV2), and Zebu Cow (*Bos indicus*, XP_070644806.1). For species with multiple isoforms, the largest isoform was used for analysis. Multiple sequence alignment was performed using Clustal Omega with default settings, and the alignment was visualized using the "serine_threonine" color scheme to highlight serine/threonine conservation. **(B)** Primate species were selected to encapsulate diverse phylogeny, including Great apes (human (*Homo sapiens*), Chimpanzee (*Pan troglodytes*), Sumatran Orangutan (*Pongo abelii*), Old world monkeys (Rhesus Macaque (*Macaca mulatta*), Olive Baboon (*Papio Anubis*), Ugandan red colobus (*Piliocolobus tephrosceles*)), and a New world monkey (common marmoset (*Callithrix jacchus*)). Identification of isoform C or D was marked if any annotated sequence from the corresponding species displayed a part of the unique amino acid sequence from C or D: SKSLSLCQINNLEKSLAAGPHHTSTHRDKPGEKQVDSNS and SECIPRWK. We note that for species where the isoform was not identified, this may be due to limitations in isoform-specific annotations in the current NCBI database.  A phylogenetic tree was created using phyloT, and isoforms were screened by manual inspection of BLASTp alignments with human isoform C and D as the query sequence.

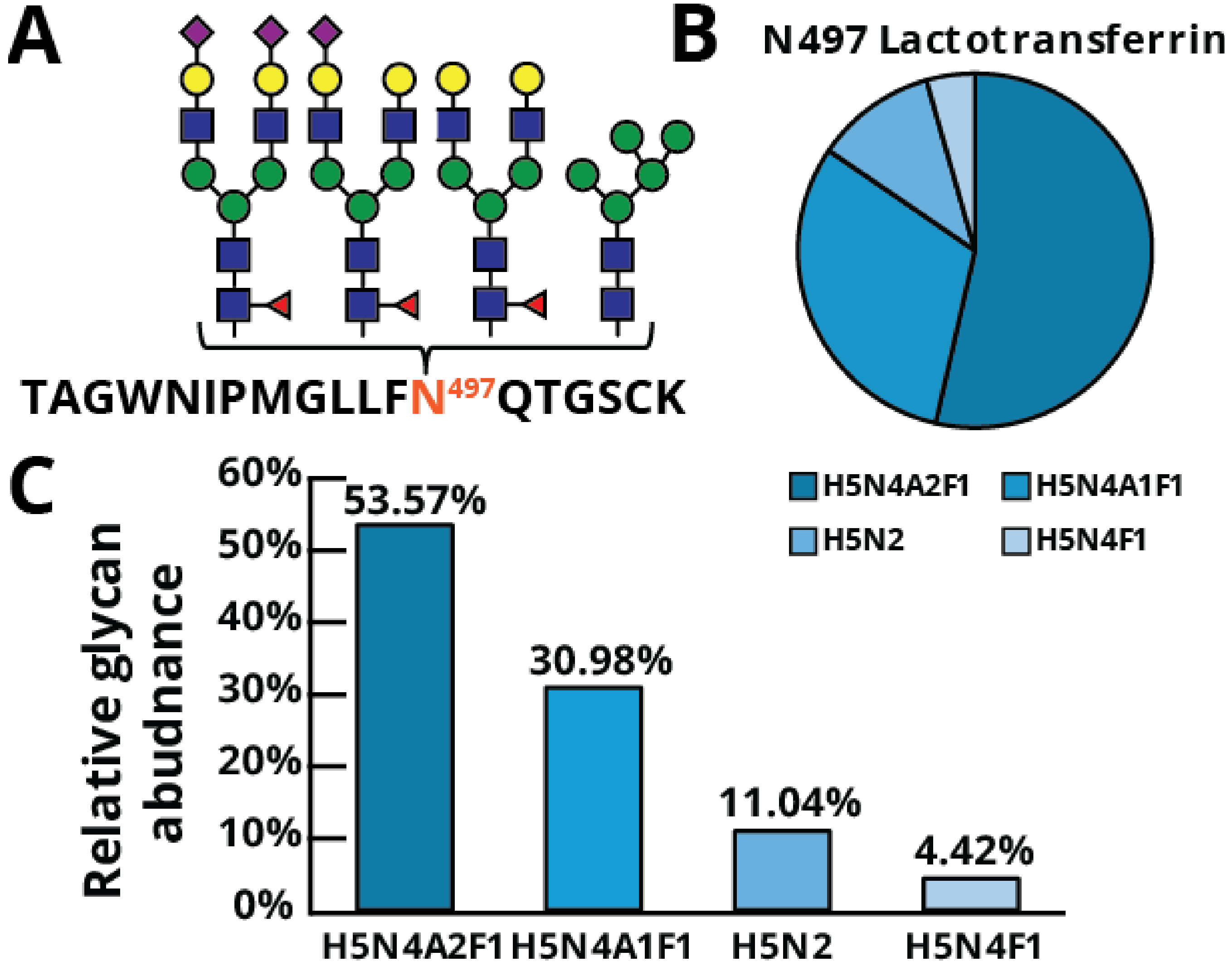


**Supplementary Figure 6. Lactotransferrin N-glycosylation analysis.**
(**A**) Lactotransferrin N-glycopeptide generated by tryptic cleavage (**B**) Relative abundances of N-glycan structures at N497 based on LFQ of AUC intensities of XICs. (**C**) Bar graph representation shown on bottom and pie chart representation shown on top right. H represents Hexose, N represents HexNAc, F represents Fucose, and A represents Neu5Ac.
